# T helper cells modulate intestinal stem cell renewal and differentiation

**DOI:** 10.1101/217133

**Authors:** Moshe Biton, Adam L. Haber, Semir Beyaz, Noga Rogel, Christopher Smillie, Karthik Shekhar, Alexandra Schnell, Zuojia Chen, Chuan Wu, Jose Ordovas-Montanes, David Alvarez, Rebecca H. Herbst, Itay Tirosh, Grace Burgin, Danielle Dionne, Michael E. Xifaras, Mei Zhang, Alex K. Shalek, Ulrich H. von Andrian, Daniel B. Graham, Orit Rozenblatt-Rosen, Hai Ning Shi, Vijay Kuchroo, Omer Yilmaz, Aviv Regev, Ramnik J. Xavier

## Abstract

In the small intestine, a cellular niche of diverse accessory cell types supports the rapid generation of mature epithelial cell types through self-renewal, proliferation, and differentiation of intestinal stem cells (ISCs). However, not much is known about interactions between immune cells and ISCs, and it is unclear if and how immune cell dynamics affect eventual ISC fate or the balance between self-renewal and differentiation. Here, we used single-cell RNA-seq (scRNA-Seq) of intestinal epithelial cells (IECs) to identify new mechanisms for ISC–immune cell interactions. Surprisingly, MHC class II (MHCII) is enriched in two distinct subsets of Lgr5^+^ crypt base columnar ISCs, which are also distinguished by higher proliferation rates. Using co-culture of T cells with intestinal organoids, cytokine stimulations, and *in vivo* mouse models, we confirm that CD4^+^ T helper (Th) cells communicate with ISCs and affect their differentiation, in a manner specific to the Th subtypes and their signature cytokines and dependent on MHCII expression by ISCs. Specific inducible knockout of MHCII in intestinal epithelial cells in mice *in vivo* results in expansion of the ISC pool. Mice lacking T cells have expanded ISC pools, whereas specific depletion of T_reg_ cells *in vivo* results in substantial reduction of ISC numbers. Our findings show that interactions between Th cells and ISCs mediated via MHCII expressed in intestinal epithelial stem cells help orchestrate tissue-wide responses to external signals.

## Introduction

The intestinal mucosa maintains a functional equilibrium with the complex luminal milieu, which is dominated by a spectrum of gut microbial species and their products. The functional balance between the epithelium and the lumen plays a central role in maintaining the normal mucosa and in the pathophysiology of many gastrointestinal disorders [1]. To maintain barrier integrity and tissue homeostasis in response to immune signals and luminal contents [1], the gut epithelium constantly regenerates by rapid proliferation and differentiation [2]. This process is initiated by intestinal stem cells (ISCs), which give rise to committed progenitors that in turn differentiate to specific IEC types [3, 4].

ISC differentiation depends on external signals from an ecosystem of non-epithelial cells in the gut niche. In particular, canonical signal transduction pathways, such as Wnt and Notch [5, 6], are essential to ISC maintenance and differentiation, and rely on signals from stromal cells [7, 8]. The intestinal tract is also densely populated by innate and adaptive immune cells, which maintain the balance between immune activation and tolerance [1, 9]. However, it is unknown if and how immune cells and the adjacent ISCs interact.

Several studies suggest an important role for immune cells in tissue homeostasis. Tissue-resident innate immune cells, such as macrophages and type 3 innate lymphoid cells (ILC3s), can play a role in regeneration of the gut [7, 10] and other tissues [11, 12]. Among adaptive immune cells, recent studies have implicated T regulatory cells (T_regs_) in regeneration within muscles, lungs, and the central nervous system [13-15]. Skin-resident T_regs_ were very recently shown to be involved in maintaining hair follicle stem cell (HFSC) renewal through Jagged1-mediated Notch signaling [16]. In the gut, mouse models of intestinal infection, T cell depletion, and inflammatory bowel disease (IBD) all display aberrant epithelial cell composition, such as goblet cell hypoplasia or tuft cell expansion [17-20]. These phenotypes have been primarily interpreted as reflecting intestinal epithelial cell dysfunction and changes in gut microbial populations [9, 17, 21, 22].

Here, we set out to identify and characterize novel mechanisms for interaction between immune cells and ISCs. Using scRNA-seq, we identified a putative molecular mechanism for CD4^+^ T cell interaction with specific subsets of Lgr5^+^ ISCs with enriched expression of MHC class II (MHCII) molecules and higher proliferation rates. We characterized this putative interaction using scRNA-Seq and *in situ* analysis of canonical *in vivo* infection models, organoid assays, and T cell-depleted, T_reg_-depleted, and inducible epithelial-specific MHCII-KO mouse models. We found that CD4^+^ T helper cells influence ISC renewal and epithelial differentiation via MHCII interaction. Our study underscores the important anatomic positioning of CD4^+^ T cell–ISC interactions in the context of ISC renewal or contraction, gut inflammation, and tumorigenesis.

## Results

### High expression of MHCII genes by ISC subsets

To identify potential mechanisms for ISC–immune cell interactions, we searched for genes that are specifically expressed by ISCs compared to other gut epithelial cells and that encode cell surface or secreted proteins capable of interacting with cognate molecules on immune cells. We collected full-length, high-coverage scRNA-seq (scRNA-seq) data of 1,522 EpCAM^+^ intestinal epithelial cells (IECs) [18] from crypt-enriched small intestine of WT and Lgr5-GFP mouse models [23] (**Methods**). Using unsupervised clustering (*k*-nearest neighbor (*k*-NN) graph-based clustering, **Methods**) of the 1,522 cells (**Table S1**) we identified 637 Lgr5-high (Lgr5^High^) stem cells (**Figure S1A,B**), as well as clusters corresponding to mature enterocytes, Paneth, goblet, tuft, and enteroendocrine cells [18]. Clustering of only the 637 ISCs (**Methods**) further partitioned the ISCs into three distinct subsets (ISC-I, -II and -III, **Figure 1A,B**), all of which express known stem cell markers [3] including *Lgr5* (**Figure S1C**). This was consistent with recent scRNA-seq reports [24, 25]. We confirmed that all three subsets comprise Lgr5^+^ ISCs using the Lgr5-GFP mouse model [26]: the three stem cell populations were strongly enriched for GFP^high^ cells (**Figure S1D**), over 90% of the GFP^high^ cells were allocated to one of the three stem cell subsets (**Figure S1E**), and the three subsets are present in similar proportions in the duodenum, jejunum, and ileum (**Figure S1F** and **Methods**). Lastly, we identified differentially expressed genes between the three subsets, as well as between all ISCs and the other IECs that are annotated as receptors or ligands for cell-cell interactions [27] (**Figure S1G** and **Table S2**).

**Figure 1.**
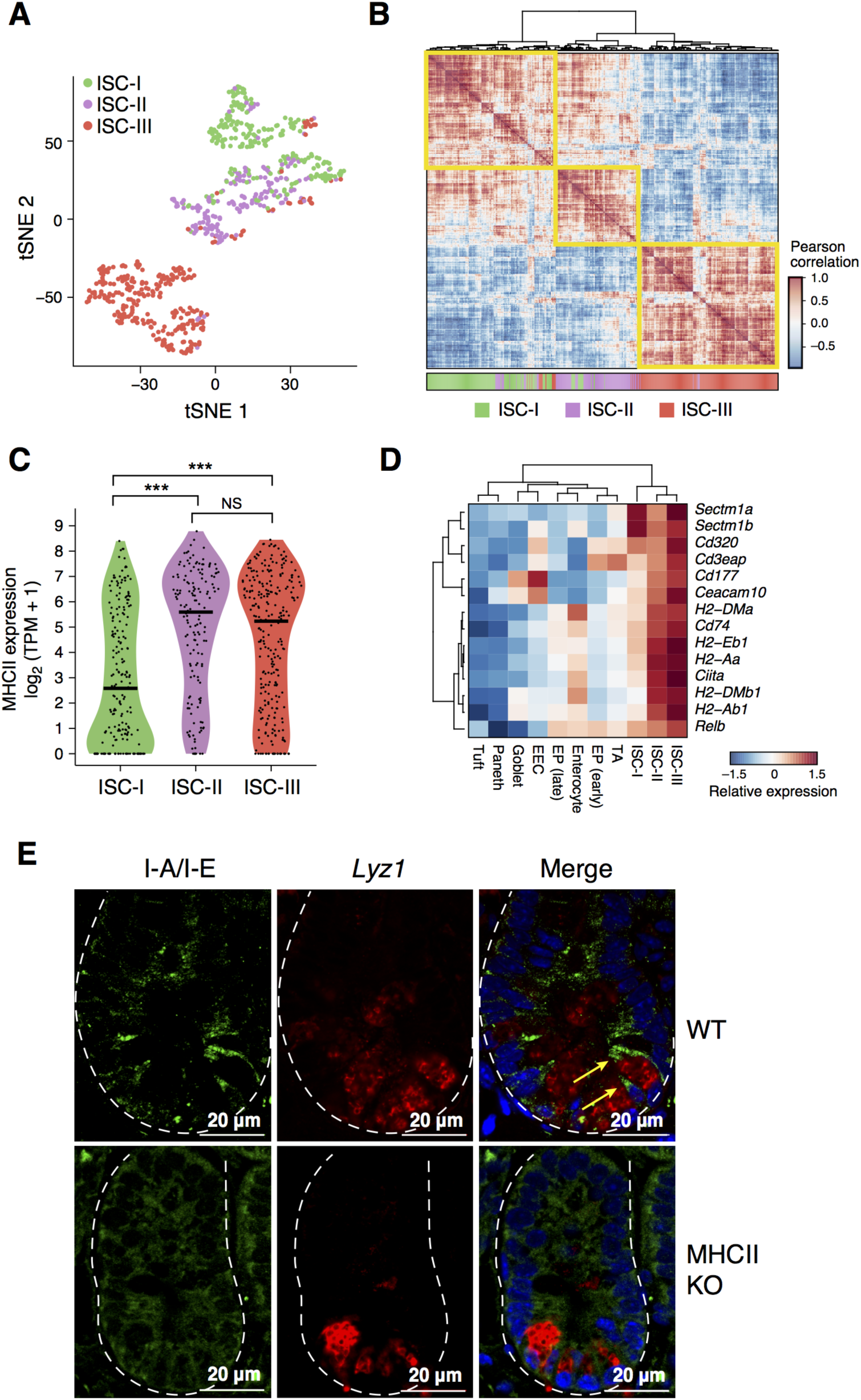
Single-cell RNA-seq reveals MHCII expression in subsets of *Lgr5*^+^ intestinal stem cells. **A,B.** Three subsets of intestinal stem cells (ISCs). Shown are a *t*-distributed stochastic neighbor (tSNE) embedding (**A**) and correlation matrix (**B**) of 637 intestinal stem cells identified by unsupervised clustering from 1,522 full-length scRNA-seq profiles (**Figure S1A, Methods**). Individual points in the tSNE embedding (**A**) correspond to single cells colored by their assignment based on *k*NN-graph-clustering (**Methods**) and *post-hoc* annotation (legend, top left). Heatmap (**B)** shows the Pearson correlation coefficient (*r*, color bar) between scores of individual cells (rows and columns) along the first 10 principal components (PCs). Color code marks ISC subsets, (bottom, **Methods**) **C.** MHCII expression in Lgr5^+^ ISCs. Violin plot of the distribution of the mean expression levels (log_2_(TPM+1), *y*-axis) of MHCII genes (*H2-Ab1, H2-Aa, Ciita, Cd74, H2-DMa, H2-DMb1*) in each of the three ISC groups. **D.** Antigen presentation genes are enriched in Lgr5^+^ ISCs. Heatmap shows the relative mean expression (row-wise Z-score of log_2_(TPM+1) values, color bar) of MHCII-related genes (rows) in each of the IEC types (columns) as defined by clustering of the 1,522 full-length scRNA-seq profiles. EP: Enterocyte progenitor, EEC: enteroendocrine cell. **E.** Validation of MHCII expression by ISCs. IFA of MHCII (I-A/I-E; green) co-stained with Paneth cell marker, Lyz1 (red) within the intestinal crypt of a wild type (WT, top row) and MHCII constitutive knockout (MHCII KO, bottom row) mouse. Yellow arrows: MHCII-expressing ISCs adjacent to Lyz1^+^ Paneth cells at the bottom of the crypt. Scale bar, 20µm.

We found that *CD74,* the invariant chain of the MHCII complex, was highly expressed and enriched in ISCs (**Figure 1C-E** and **Figure S1G**). Moreover, other MHCII genes were among the most strongly expressed by two out of three ISC subsets (ISC-II and -III) (**Figure 1C,D**) compared to other IECs (**Figure S2A**). These included many canonical components of the MHCII machinery, including *H2-Ab1*, *H2-DMb1*, *H2-DMa*, *H2-Aa*, *Cd74*, and the recently discovered co-stimulatory molecules *Sectm1a* and *Sectm1b* [28] (**Figure 1D**), but not the canonical co-stimulatory molecules CD80 and CD86 (**data not shown**). Although MHCII expression has been previously reported in intestinal epithelial cells [29-32], it was not shown to be enriched in ISCs. We found that MHCII expression in the ISC^MHCII+^ (ISC-II and -III) groups was the highest among all IECs at both the mRNA and protein levels (**Figure S2A,B**). We confirmed MHCII protein expression by ISCs using an immunofluorescence assay (IFA) and immunohistochemistry (IHC) with anti-MHCII antibodies in wild type mice, and its absence in a constitutive MHCII knockout (KO) [33] (**Figure 1E** and **Figure S2C**).

### MHCII-expressing ISCs are more proliferative

The three ISC subsets vary not only in their expression of the MHCII system, but also in their expression of signatures of the cell-cycle [34, 35] (**Figure 2A,B**). The subset with highest MHCII expression consisted primarily of cells in G1/S. The second subset, with lower but significant MHCII expression, had cells spanning several phases of the cell-cycle including G2/M. We concluded that cells in both of these subsets are likely in highly proliferative states and termed these ISC-II and ISC-III, respectively. In contrast, the cells in the subset with low or no detectable expression of MHCII, termed ISC-I, also had low G1/S and G2/M scores (129 of 209 cells) and likely represented cells in G0. The low-cycling state of ISC-I was further supported by the higher expression of the histone methylase *Kdm5b*, which is highly expressed in post-mitotic differentiated cells of the small intestine (**Figure S2D,E**) and in low-cycling or quiescent cells in other systems [34, 36-39]. Such heterogeneity in the proliferative state of ISCs has been recently reported, including a quiescent ISC subset, which is enriched for *Mex3a* and correlates well with our low cycling ISC-I subset (data not shown) [24, 25]. Importantly, while the cell-cycle status aligns with the partitioning of ISC-I, II and III subsets, the ISC subsets are discernable even when we exclude canonical cell-cycle genes (**Figure 2C, Figure S3A**, **Table S3** and **Methods**) and even when analyzing only the 183 ISCs scoring at G0 (**Figure S3B**).

**Figure 2.**
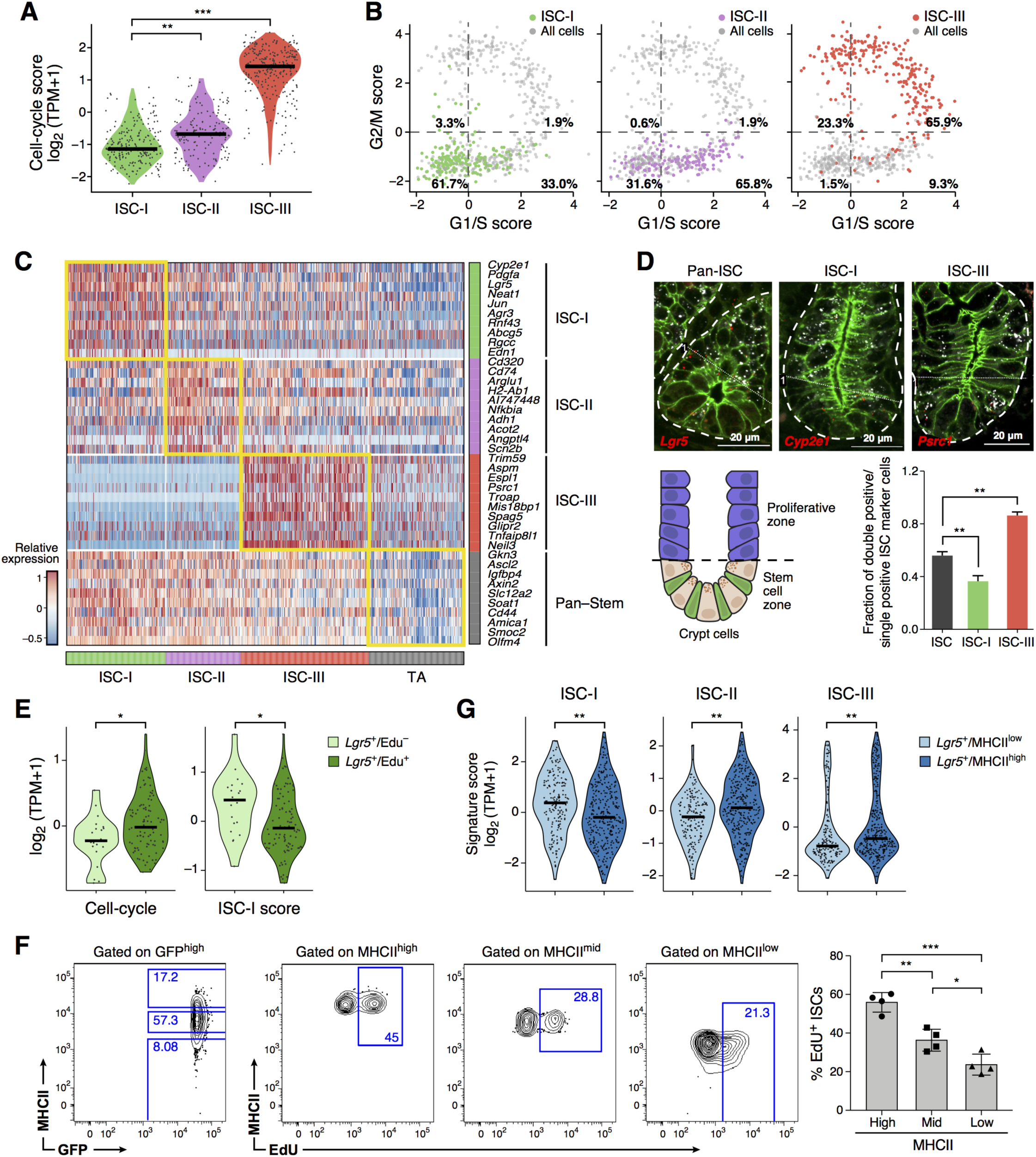
MHCII expression is correlated with ISC proliferation. **A,B.** Distinct cell-cycle characteristics in the three ISC subsets. (**A**) Violin plot shows the distribution of expression scores (*y*-axis) for a signature of cell-cycle genes (**Methods**) in each stem cell subset (*x*-axis). ** *p*<10^-5^, *** *p*<10^-7^ (Mann-Whitney U-test). (**B**) Scatter plots show the signature score for 637 ISCs (points, **Methods**) for G1/S genes (*x*-axis) and G2/M genes (*y*-axis). Cells are colored by their cluster assignment to ISC subsets. **C.** Gene signatures of ISC subsets. Heatmap shows relative expression level (row-wise Z-score of log_2_(TPM+1) values, color bar) of ten representative genes from each ISC subset signature and a pan-stem signature (rows, right color bar) in 637 individual stem cells and 201 TA progenitors (columns, bottom color bar as in **(A)**) identified from 1,522 scRNA-seq profiles. Gene signatures are identified based on our analysis as well as from a previously published signature of stem cell genes of a bulk dataset [3]. **D.** Identification of proliferating stem cells in intestinal crypts. Upper panels: Combined single-molecule fluorescence *in situ* hybridization (smFISH) with immunofluorescence (IFA) of FFPE sections of intestinal tissue from wild type mice, showing the pan-stem cell marker *Lgr5* (upper left), ISC-I marker *Cyp2e1* (upper middle) and ISC-III marker *Psrc1* (upper right) all in red and *mKi67* in white. Cell borders were assessed with E-cadherin (green); scale bar, 20μm. Bottom left: Schematic of the lower crypt fraction (‘stem cell zone’), in which co-expression of stem cell markers (*Lgr5* and *Cyp2e1* or *Psrc1*) and the proliferation marker *mKi67* was quantified. Bottom right: Bar plot showing the fraction (*y*-axis) of cells which are positive for *mKi67* out of all cells positive for each stem cell marker (lower right panel). *n*=4 mice, 10 crypts on average per mouse (** *p*<0.0025, *t*-test; error bars: SEM). **E**. Enrichment of ISC-I in EdU^-^ cells. Violin plots show the distribution of signature scores for the cell-cycle (left, signature as in **(A)**) and ISC-I (right, signature as in **(C)**) from FACS sorted EdU^-^ Lgr5^+^ ISCs (light green) or EdU^+^ Lgr5^+^ ISCs (dark green) after *in vivo* EdU labeling and profiled using single-nucleus RNA-seq (Div-Seq). **F.** Higher proliferation of ISCs with high MHCII expression. FACS plots of ISCs gated on GFP-high (Lgr5^+^, left) binned into subsets with low, intermediate and high MHCII expression (middle panels, *y*-axis), and then gated on EdU incorporation (middle panels, *x*-axis). Bar plot (right) shows the fraction (percentage, *y*-axis) of EdU^+^ cells within each MHCII expression level (*n*=4 mice, * *p*<0.05, ** *p*<0.005, *** *p*< 0.0005, *t*-test, error bars: SEM). **G**. ISC subset signatures across MHCII expression. Violin plots show the distribution of signature scores (as in (**C)**) for ISC-I, ISC-II, and ISC-III subsets (left to right), across scRNA-seq profiles from 326 Lgr5^+^ MHCII^high^ (light blue) and 177 Lgr5^+^MHCII^low^ (dark blue) cells (individual black dots). Horizontal black line denotes the median (** *p*<0.005, *** *p*< 0.0005, Mann-Whitney U-test).

We validated the association between the ISC subsets, MHC-II, and cell-cycle status by co-staining *in situ* (**Figure 2D**), by *in vivo* EdU labeling followed by single-nucleus RNA-seq (Div-Seq) [40] (**Figure 2E**), and by determining the proportion of EdU^+^ cells in subsets of GFP^high^ cells with different levels of MHCII expression (**Figure 2F**). We also sorted single MHCII^high^ and MHCII^low^ ISCs from Lgr5-GFP mice and collected 503 full-length scRNA-seq profiles. MHCII^high^ ISCs had a higher proportion of cells with high scores for ISC-II and -III state signatures, whereas MHCII^low^ ISCs had a higher proportion of cells scoring highly for the ISC-I state, consistent with our *in silico* analysis (**Figure 2G** and **Figure S3C**). Taken together, these results support an association between MHCII expression and proliferative state within ISCs *in vivo*. Thus, the heterogeneity of Lgr5^+^ stem cells and MHC II expression is correlated with proliferation rates.

### T helper subsets and their signature cytokines regulate ISC renewal in organoids

We hypothesized that ISCs may interact with CD4^+^ T helper (Th) cells via MHCII recognition and, as a consequence, CD4^+^ Th cells may affect ISC fate via cytokine–receptor interaction. Importantly, IECs, including ISCs, express receptors for Th cytokines interferon gamma (IFNγ), interleukin-10 (IL-10), IL-13, IL-4 and IL-17A (data not shown). Furthermore, intra-vital imaging showed that CD4^+^ Th cells can be in very close proximity to stem cells in small intestinal crypts (**Figure S4A** and **movie S1**). Moreover, scRNA-seq of IECs following infection of mice with *Salmonella enterica* (*Salmonella*) or *Heligmosomoides polygyrus* (*H. polygyrus)* [18], which induce Th1 (**Figure S4B-D** and **Table S4**) and Th2 responses, respectively, shows not only distinct shifts in the proportions of post-mitotic cells, such as tuft (in *H. polygyrus*) and Paneth (in *Salmonella*) cells [18] (**Figure S4E-F**), but also a reduction in ISC and stemness programs, and especially in ISC-I cells (**Figure S4G-J**). The observed elevation in ISC^MHCII+^ programs is consistent with our hypothesis, but in principle many other indirect cellular and molecular mechanisms in the complex cellular ecosystem of the crypt may also be involved.

To dissect the potential interactions between T helper cells and ISCs independently of other contributions to the niche, we therefore next used the intestinal organoid system [26] in which immune cells are natively absent but can be added in a controlled manner [41]. We introduced either specific CD4^+^ T helper subsets (**Figure S5A**) or their corresponding signature cytokines to organoid cultures, and used scRNA-seq to identify changes in the proportions or expression programs of ISCs. In one set of experiments, we co-cultured organoids with CD4^+^ T cells that were polarized *ex vivo* towards Th1, Th2, Th17, and iT_reg_ cells [42] (**Figure S5B**). In a parallel set, we stimulated organoids derived from C57BL/6J WT mice with key cytokines produced by each of the four T helper subsets: IFNγ (Th1), IL-13 (Th2), IL-17a (Th17), and IL-10 (inducible T_reg_, iT_reg_). In each experiment, we collected droplet-based scRNA-seq profiles (**Methods**). For co-cultures, we computationally distinguished (*post-hoc*) T cells from epithelial cells by their profiles (**Methods**) and confirmed the Th cell state by mRNA expression of signature cytokines and transcription factors (**Figure S5C**). Although *ex vivo* polarized T helper cells share many hallmarks with their *in vivo* counterparts, they do not perfectly recapitulate them. In particular, Th2 differentiation yielded only 16.5% IL-4 and IL-13 expressing cells, while other T helper subsets had higher differentiation rates (**Figure S5B**). There are also several differences between organoids and *in vivo* IECs (**Figure S5D-G**): Organoids are enriched for stem cells [26, 43] (**Figure S5D**), the goblet and Paneth lineages do not fully diverge (**Figure S5E**), also in independently-generated organoids [44] (**Figure S5F**), and MHCII expression was not detected in our organoid culture (**Figure S5G**); thus, any impact of Th cell co-cultures is likely mediated through cytokine secretion from the polarized Th cells.

Each of the Th co-cultures or corresponding cytokine treatments resulted in a distinct modulation of the organoid ISC compartment (**Figure 3A,B** and **Figure S6A-C**). Strikingly, co-cultures with iT_regs_ and treatment of organoids with their associated cytokine IL-10 led to organoid ISC expansion (**Figure 3A,B, Figure S6A-C, Methods** and **Table S5**), while co-cultures with Th1, Th2 and Th17 cells or treatment with IL-13 or IL-17 all reduced the size of the ISC pool of the organoids. Consistent with their depleted stem cell pool, organoids co-cultured with Th1, Th2, or Th17 cells or treated with IL-17a all showed elevated numbers of TA cells (*p*<10^-4^, hypergeometric test, **Figure 3A,B**). Note, for IFNγ, we used a low concentration (0.5u/ml) to avoid organoid apoptosis [45], which did not elicit any effect, while Th1 co-culture resulted in elevation of MHCII signature in IECs (**Figure 3A,** top and **Figure S5G**). In addition, the treatments impacted cell differentiation: IL-13 treatment decreased the proportion of secretory ‘Paneth-goblet’ cells, and increased tuft cells (**Figure 3A,C**) [19, 46]; Th1 co-culture up-regulated Paneth cell-specific genes (**Figure S6D-F**), consistent with *in vivo* observations (**Figure S4F**); and Th2 cell co-cultures had the opposite effect (**Figure S6D,E**).

**Figure 3.**
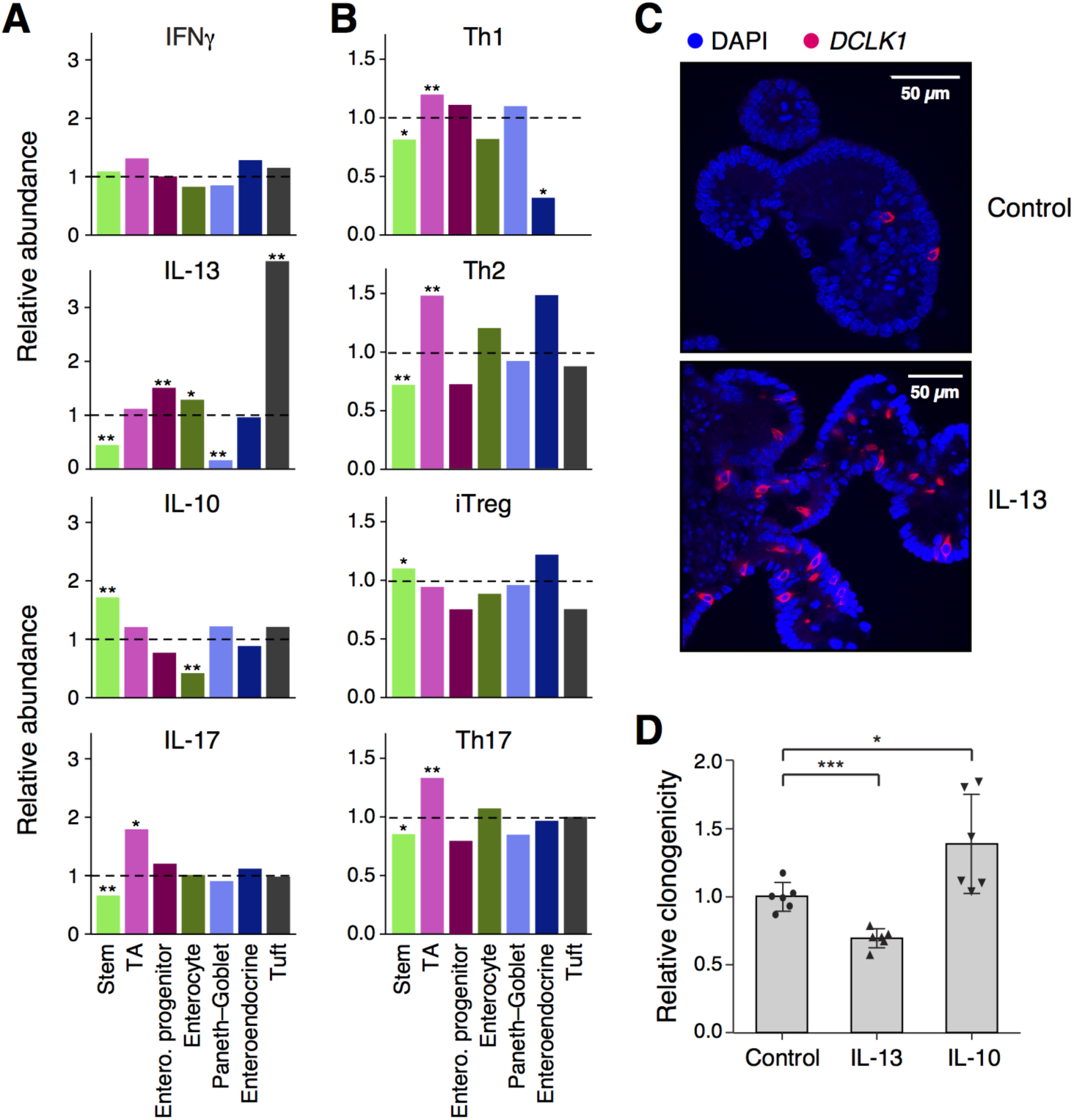
T helper cells and their key cytokines impact ISC number and differentiation in intestinal organoids. **A,B.** Shifts in cell type composition within organoids treated with CD4^+^ Th signature cytokines (**A**) or co-cultured with Th cells (**B**). relative abundance (*y*-axis) of each IEC-type under each condition relative to their proportions in control organoids (dashed line). Subsets were identified by unsupervised clustering of 23,177 single cells obtained from the Th co-culture and cytokine conditions and annotated *post-hoc* (**Figure S5D, Methods**). * *p* <0.01, ** *p* <10^-4^ (hypergeometric test, **Methods**) **C.** DCLK1^+^ tuft cell expansion following IL-13 treatment. IFA of DCLK1 (red) in control organoids (top) and IL-13-treated organoids (bottom). Scale bar, 50µm. **D.** Altered clonogenicity following pre-treatment with different cytokines. Bar plot shows the relative clonogenicity of organoid cultures (*y*-axis, relative to the mean value of control organoids) defined by the number of organoids in cultures re-seeded after treatment with IL-10 or IL-13 (*x*-axis). Dots: technical replicates. Error bars: SD, * *p*<0.05, *** *p*<0.0005, *t*-test.

The effects of Th cell subsets and cytokines on ISC numbers suggest that they affect ISC renewal potential, which in turn should affect the ability of ISCs to form organoid cultures. To test this hypothesis, we assessed whether key cytokines affect ISC clonogenicity [47]. We reseeded equal numbers of cytokine-treated organoids in new cultures and quantified the number of organoids after three days (*n*=6 replicates per each group, **Methods**). Consistent with our hypothesis, there was a significant reduction in the clonogenicity of organoids treated with the ISC-reducing cytokine IL-13, whereas the ISC-expanding cytokine IL-10 induced higher clonogenicity (**Figure 3D**), confirming the ability of this T_reg_-generated cytokine to rejuvenate the stem cell pool.

### Elevation in ISC pool under epithelial MHCII ablation *in vivo*

Since the MHCII system is not expressed in organoids, we next assessed its role in IECs *in vivo* by its conditional KO. We crossed H2-Ab1^fl/fl^ [48] to Villin-Cre-ER^T2^ [49] mice, generating a mouse model of specific and inducible MHCII knockout in IECs (MHCII^Δgut^). We profiled 1,559 IECs from the MHCII^Δgut^ mice (*n*=5) 10 days after Tamoxifen induction and 1,617 IECs from floxed control (MHCII^fl/fl^) littermates (*n*=5 mice). We validated that MHCII is successfully knocked-out in EpCAM^+^ IECs (**Figure 4A** and **Figure S7A**), but not in CD11b^+^ dendritic cells in the mesenteric lymph node (**Figure S7B**).

**Figure 4.**
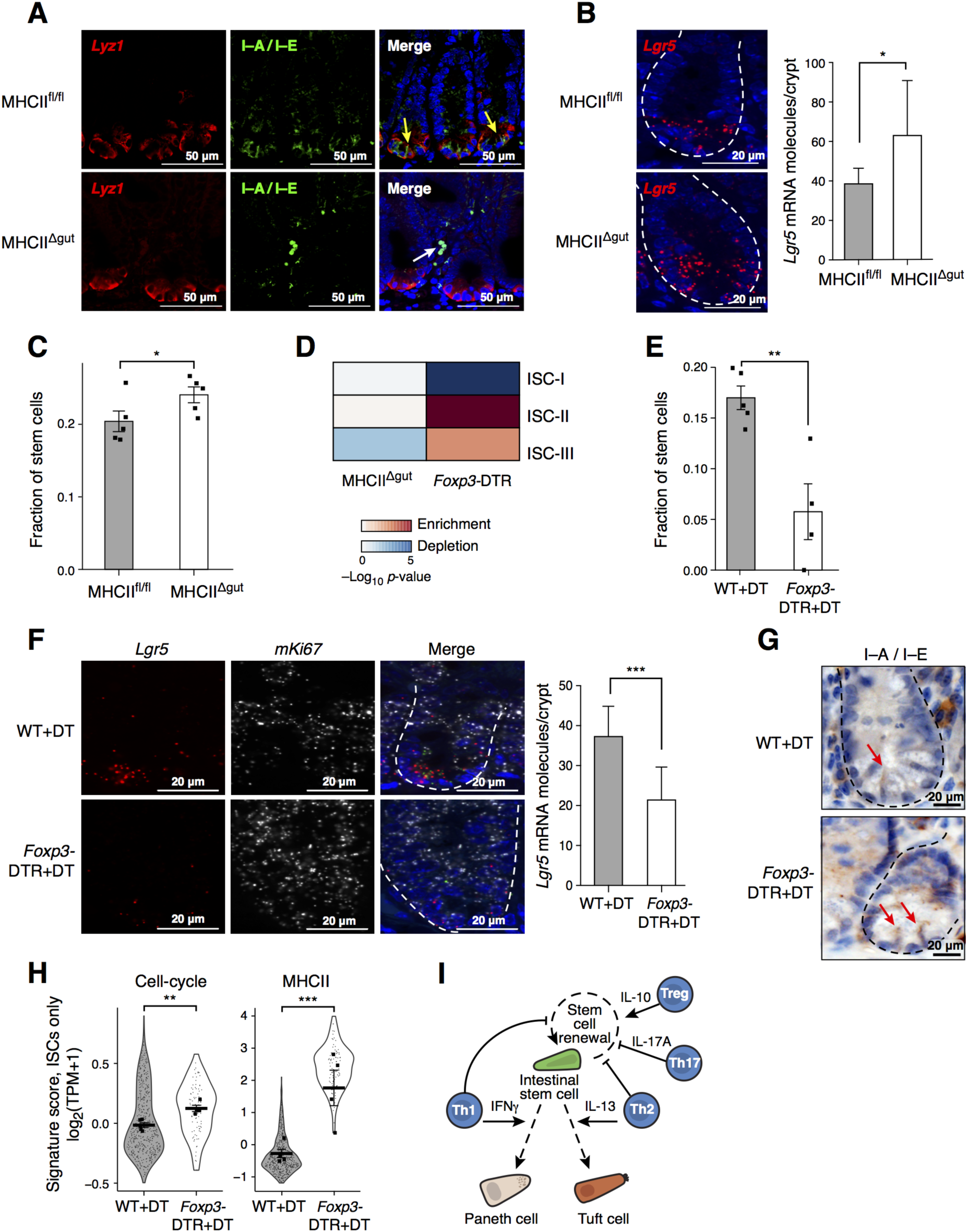
MHCII depletion leads to increased ISC numbers while T_reg_ ablation results in reduced ISC pool. **A-D.** Expansion of the ISC pool following epithelial specific ablation of MHCII. (**A**) Validation of epithelial-specific MHCII-KO (MHCII^Δgut^) mouse. IFA of Lyz1 (red) and MHCII (I-A/I-E, green) in MHCII^fl/fl^ (top row) and MHCII^Δgut^ mice (bottom row). Yellow arrow: MHCII^+^ epithelial cell, white arrow: MHCII^+^ non-epithelial cell. (**B**) smFISH of the expression of *Lgr5* (red) within intestinal crypts from MHCII^fl/fl^ controls (*n*=5, top left) and MHCII^Δgut^ mice (*n*=5, bottom left). Bar plot (right) shows the number of *Lgr5* mRNA molecules per crypt (*y*-axis) in MHCII^fl/fl^ and MHCII^Δgut^ mice (*x*-axis). *n*=2 mice and 8 fields per group. Error bars: SD (* *p*<0.05, *t*-test). (**C**) Bar plot shows the fraction of ISCs within EpCAM^+^ cells (*y*-axis, determined by unsupervised clustering) in MHCII^fl/fl^ and MHCII^Δgut^ mice (points, *x*-axis). Error bars are SEM. (* FDR<0.05, likelihood-ratio test, **Methods**). (**D**) Heatmap shows the significance of changes in signature scores (-log_10_(*p*-value), Mann-Whitney U-test, of enrichment (red) and depletion (blue), **Methods**) of MHCII^Δgut^ or T_reg_ ablation compared to control mice (columns) for signatures (rows) associated with the three ISC subsets. **E-H.** Reduction in ISC numbers and shifts toward ISC-II and ISC-III states in T_reg_ depleted mouse model. (**E**) Bar plot shows the fraction of ISCs within EpCAM^+^ cells (*y*-axis) detected by unsupervised clustering in WT and *Foxp3*-DTR mice both treated with DT). Error bars are SEM. (** FDR<0.005, likelihood-ratio test, **Methods**). (**F**) smFISH of FFPE sections of intestinal tissue from wild type mice (top) or *Foxp3*-DTR mice (bottom) both treated with DT, showing from left to right, the stem cell marker *Lgr5* (red), *mKi67* (white) and a merge; scale bar, 20μm. Bar plot (right) shows the number of *Lgr5* mRNA molecules per crypt (*y*-axis) in WT and *Foxp3*-DTR mice treated with DT (*x*-axis). *n*=2 mice and 8 fields per group, error bars are SD (*** *p*<0.001, *t*-test). (**G**) Immunohistochemistry (IHC) of MHCII (I-A/I-E; brown) co-stained with hematoxylin (blue) within the intestinal crypt of wild type (top) and *Foxp3*-DTR (bottom) mice both treated with DT. Red arrows indicate MHCII^+^ ISCs. (**H**) A shift towards proliferative ISC states following T_reg_ depletion. Violin plots of the distribution of scores (**Methods**) for the cell-cycle (left; as in **Figure 2**) and MHCII genes (right; as in **Figure 1**) in ISCs (small points) from WT (*n*=5; 2,572 cells) and *Foxp3*-DTR (*n*=4, 815 cells) mice. Squares: mean score per mouse; thick bar: overall mean; error bars: SEM. (* *p*<0.05, ** *p*<0.005, *** *p*<5×10^-4^, likelihood-ratio test). **I.** Proposed model of a novel interaction between CD4^+^ T helper cells and ISCs mediated by MHCII. T helper cell subsets (blue nodes) modulate (solid arrows) the differentiation (dashed arrows) of ISCs (green). T_regs_ and their key cytokine IL-10 promote stem cell renewal, while Th17 cells and their cytokine IL-17a reduce stem cell renewal and promote differentiation. Both Th1 and Th2 suppress stem cell renewal and promote specific differentiation towards Paneth cells (tan) and tuft cells (orange), respectively.

Strikingly, the fraction of *Lgr5*^+^ cells was 31.3% higher in MHCII^Δgut^ mice (*p*<0.05, likelihood-ratio test, **Figure S8A**), which we confirmed by *Lgr5*-smFISH (**Figure 4B**), and the proportion of ISCs as defined by unsupervised clustering (**Figure S8B,C**) which was 17.6% higher (FDR<0.05, likelihood-ratio test, **Figure 4C**). Consistently, stem cell markers are overrepresented (*p*<10^-6^, hypergeometric test, **Figure S8D**) among the genes up-regulated in the MHCII^Δgut^ (FDR<0.05, likelihood-ratio test), including canonical ISC markers (*e.g.*, *Lgr5*, *Olfm4*, *Smoc2*, and *Igfbp4,* **Figure S8D**). Furthermore, we separately analyzed only MHCII^Δgut^ ISCs in which *H2-Ab1* is confirmed to be silenced (defined as no detectable mRNA) or only MHCII^Δgut^ ISCs in which *H2-Ab1* mRNA is still expressed (**Figure S8E**, bottom left *vs*. right). We find that in MHCII^Δgut^ ISCs in which *H2-Ab1* is confirmed to be silenced, expression of stem cell markers [3] was significantly higher than in stem cells still expressing *H2-Ab1* (*p* < 0.05, likelihood-ratio test). Finally, the ISC-III signature score was significantly lower in stem cells from MHCII^Δgut^ mice (**Figure 4D**), suggesting that the ISCs in the expanded pool are shifted toward the ISC^MHCII-^ state. Taken together, these data suggest that MHCII^Δgut^ increased ISC numbers and the expression of stem cell markers.

### T cells modulate ISC renewal and differentiation *in vivo*

Our organoid assays predicted that Th cell subsets have distinct effects on intestinal epithelial cell differentiation. To demonstrate the relevance of the T cell-ISC interaction *in vivo*, we first assessed two T cell-deficient mouse models. First, we profiled 2,967 individual IECs isolated from athymic B6 nude mice [50] (*n*=2), characterized by T cell depletion. Unsupervised clustering revealed a markedly higher fraction of stem cells (52.5% increase, FDR<10^-3^, likelihood-ratio test, **Methods**) compared to control mice (*n*=6, **Figure S9A,B,E**). Consistently, stem cell markers were enriched (56 of 1,804 genes, *p*<10^-6^, hypergeometric test, **Figure S9A** and **Table S6**) among genes overall up-regulated in cells of nude *vs*. controls (FDR < 0.05, likelihood-ratio test). Similar analysis of 9,488 individual IECs profiled from TCRβ-KO mice (*n*=2) [51], characterized by a lack of α/β T cells, also showed a significant expansion of the ISC pool (35.0% increase, FDR < 0.05, likelihood-ratio test; **Figure S9C-E**). We confirmed the increased ISC numbers *in situ* in both T cell depleted models using *Lgr5* single-molecule FISH (smFISH, **Figure S9F**).

### T_reg_ cells are essential to maintain the ISC niche *in vivo*

Our organoid assays also predicted that T_reg_ cells promote renewal of the ISC pool. To test for this effect *in vivo*, we used the *Foxp3*-DTR mouse [52], in which T_reg_ cells are specifically depleted upon application of diphtheria toxin (DT). We profiled 3,387 IECs from both *Foxp3*-DTR (*n*=4) and matched control mice (*n*=5) treated with DT for 7 days and confirmed T_reg_ ablation in the lamina propria (**Figure S10A**). At this time point, there were no signs of increased cell death in the small intestinal crypts or of major tissue inflammation in IECs of *Foxp3*-DTR *vs*. control mice (**Figure S10B**), suggesting that the longer term effects of T_reg_ depletion are not yet apparent [52]. However, consistent with our hypothesis, there was a substantial reduction (66.3% decrease, FDR<0.005, likelihood-ratio test) in ISC numbers in the epithelia of the *Foxp3*-DTR mice, as assessed by unsupervised clustering (**Figure 4E**), the fraction of cells in which *Lgr5* mRNA was detected (*p*<0.005, likelihood-ratio test, **Figure S10C**), and smFISH (**Figure 4F**). Consistently, stem cell marker genes were overrepresented among those down-regulated across all cells in T_reg_-depleted mice (*p*<0.005, hypergeometric test, **Figure S10D**). There was also a substantial depletion of mature enterocytes (5.8-fold decrease, FDR<10^-5^), and expansion of tuft cells (4.1-fold increase, FDR<0.005) (**Figure S10E**), which we confirmed by IFA staining (**Figure S10F**). We did not observe significant changes in the expression of Notch signaling pathway components (*p*=1), or Notch targets [53] (*p*=0.31, hypergeometric test), which a recent study implicated in regulation of hair follicle stem cells by T_regs_ [16] (data not shown).

All cell types in the *Foxp3*-DTR mice, including ISCs, showed strongly elevated expression of MHCII genes (*p*<5×10^-4^, likelihood-ratio test, **Figure 4G**). Amongst stem cells, there was an increase in proliferation, as indicated by both the distribution of cell-cycle signature scores and mKi67 staining (*p*<0.005, likelihood-ratio test, **Figure 4F** and **Figure S10B**). Furthermore, and also consistent with our predictions, ISCs from T_reg_-ablated *Foxp3*-DTR mice had an increased proportion of MHCII positive, proliferative ISCs and a decrease in ISC^MHCII-^ (ISC-I, **Figure 4D**).

## Discussion

Previous studies of stem cell dynamics and differentiation processes [54, 55], focused on the role of the epithelial-intrinsic or stromal niche signals using lineage tracing. Here, we investigated the possibility of interactions between adaptive immune cells and ISCs. Combining scRNA-seq with homeostatic or perturbed conditions that manipulate either T helper cells, their cytokines, or MHCII expression by epithelial cells allowed us to assay comprehensive “snapshots” of ISC abundance and the fate of their progeny, followed by *in silico* inference of cell states and differentiation. In unperturbed mice, the expression of MHCII is high yet variable across ISCs, such that both ISC-II and III (ISC^MHCII+^) express high levels of the MHCII molecules, whereas ISC-I (ISC^MHCII-^) do not. Using controlled manipulation experiments in organoids and mice followed by scRNA-seq, we established a crosstalk between Th cells and ISCs.

In particular, our *in vitro* and *in vivo* results support a model in which Th cells interact with ISCs via MHCII molecules, impacting the ISC pool and resultant differentiation pathways through their key cytokines (**Figure 4I**). In this model, T_reg_ cells, which are enriched in the small intestine, maintain the ISC niche. They may be elevated after a strong inflammatory response [56] to serve as a feedback effectors in order to replenish and maintain stem cell numbers. Conversely, Th1 and Th2 cells or their signature cytokines both reduce ISC numbers, and bias IEC differentiation toward specific epithelial cell-types, perhaps in order to respond to either bacterial (Th1 cells, Paneth cell increase) or parasitic (Th2 cells, tuft cell increase) insults. Th17 cells, which are highly enriched in the small intestine [57], reduce the number of ISCs, which may reflect a shift in the balance between stem cell renewal and differentiation. In this way, epithelial and immune response could be integrated to titrate responses to dense luminal flora, avoiding continuous inflammation, while reacting to pathogens: first, the intestinal stem cells utilize the equilibrium of pro-inflammatory and anti-inflammatory signals to balance between renewal and differentiation; second, distinct Th cell subsets can boost the desired immune response by affecting renewal and differentiation processes of the gut epithelia concordantly with signals arriving from the gut lumen. If this novel role for MHCII in T cell communication with stem cells also exists in other mucosal or non-mucosal compartments, it may open the possibility of a general mechanism in which adaptive immune cells regulate parenchymal stem cells in order to maintain tissue homeostasis under normal and pathological conditions.

## Acknowledgements

We thank Leslie Gaffney for help with figure preparation, the Broad Flow Cytometry Facility: Patricia Rogers, Stephanie Saldi and Chelsea Otis and Tim Tickle for help with the Single Cell Portal. Work was supported by the Klarman Cell Observatory at the Broad Institute (AR and RJX), HHMI (AR), and a Broad*next*10 award (AR an RJX). MB was supported by a postdoctoral fellowship from the Human Frontiers Science Program (HFSP). RJX funded by DK043351, DK097485 and Helmsley Charitable trust. ASK in a member of Searle Scholars Program, the Beckman Young Investigator Program and the recipient of the NIH New Innovator Award DP2 OD020839. ÖHY is funded by CA211184 and AG045144 NIH grants and is the recipient of the Sidney Kimmel Scholar Award, the Pew-Stewart Trust Scholar Award and a member in the American Federation of Aging Research. AR is a member of the SAB for ThermoFisher Scientific, Syros Pharmaceuticals, and Driver Group.

## Supplementary Materials for T helper cells modulate intestinal stem cell renewal and differentiation

### Materials and Methods

#### Experimental Methods

##### Mice

All mouse work was performed in accordance with the Institutional Animal Care and Use Committees (IACUC) and relevant guidelines at the Broad Institute and MIT, with protocols 0055-05-15 and 0612-058-15, respectively. Seven to ten weeks old female or male C57BL/6J wild-type, Lgr5-EGFP-IRES-CreER^T2^ (Lgr5-GFP), MHCII-KO, Foxp3-DTR, B6 Nude and TCRβ-KO mice were obtained from the Jackson Laboratory (Bar Harbor, ME), while MHCII-EGFP mice were obtained from Hidde Ploegh’s lab. All mice were housed under specific-pathogen-free (SPF) conditions at either the Broad Institute or MIT animal facilities; infection experiments were conducted at the laboratory of Dr. HN Shi, maintained under specific pathogen-free conditions at Massachusetts General Hospital (Charlestown, MA), with protocol 2003N000158.

*BrdU and EDU incorporation:* EdU was injected intraperitoneally (IP) into Lgr5-GFP mice at 100 mg kg^-1^ for 2 or 4 hours before tissue collection.

*S. enterica and H. polygyrus infection.* C57BL/6J mice (Jackson Laboratory) were infected with 200 third-stage larvae of *H. polygyrus* or 10^8^ *S. enterica*. *H. polygyrus* was propagated as previously described [58]. Mice were sacrificed 3 and 10 days after *H. polygyrus* infection. For the MHCII blocking experiment, mice infected with *H. polygyrus* were injected with 500µg of blocking anti-mouse MHCII antibody (BioXCell) or Rat IgG2b isotype control (BioXCell) one-day prior to and for 2 consecutive days after *H. polygyrus* infection. For *S. enterica* infection, mice were infected with a naturally streptomycin-resistant SL1344 strain of *S. Typhimurium* (10^8^ cells) as previously described [59] and were sacrificed 48 hours after infection.

*Foxp3-DTR.* Foxp3 and wild-type C57BL/6J mice were injected intraperitoneally with diphtheria toxin (DT) at 22.5ng/g body weight every other day for one week and then sacrificed.

*MHCII deletion in intestinal epithelial cells.* Cre activity was induced in 7-10 weeks old mice by intraperitoneal injection (IP) of Tamoxifen (SIGMA), diluted in corn oil, 4 mg per injection, 3 times, every other day. Mice were sacrificed 10 days after the first injection.

##### Cell dissociation and crypt isolation

*Crypt isolation*. For all mice, crypts were isolated from the whole small intestine or the duodenum, jejunum and ileum compartments to account for regional distribution of Lgr5^+^ stem cells. The small intestine was extracted and rinsed in cold PBS. The tissue was opened longitudinally and sliced into small fragments roughly 0.2 cm long. The tissue was incubated in 20mM EDTA-PBS on ice for 90 min, while shaking every 30 min. The tissue was then shaken vigorously and the supernatant was collected as fraction 1 in a new conical tube. The tissue was incubated in fresh EDTA-PBS and a new fraction was collected every 30 min. Fractions were collected until the supernatant consisted almost entirely of crypts. The final fraction (enriched for crypts) was washed twice in PBS, centrifuged at 300g for 3 min, and dissociated with TrypLE Express (Invitrogen) for 1 min at 37ºC. The single-cell suspension was then passed through a 40µm filter and stained for fluorescence-activated cell sorting (FACS) machine (Astrios) sorting for either scRNA-seq method (below).

*Immune cell isolation.* Immune cells from the Lamina Propria were isolated enzymatically by incubating the small intestine with Liberase TM (100 μg/mL, Sigma) and DNaseI (100 μg/mL, Sigma) for 30 min at 37ºC. Immune cells were also isolated from the mesenteric lymph nodes (mLN). Cells were then incubated with CD3, CD4, CD45, or CD11b FACS-labeled antibodies and sorted for scRNA-seq.

##### Cell sorting

*Plate-based, full-length scRNA-seq*. FACS (Astrios) was used to sort one single cell into each well of a 96-well PCR plate containing 5µl of TCL buffer with 1% 2-mercaptoethanol. The cells were stained for 7AAD^-^ (Life Technologies), CD45^-^ (eBioscience), CD31^-^ (eBioscience), Ter119^-^ (eBioscience), EpCAM^+^ (eBioscience), and for specific epithelial cells were also stained for CD24^+/-^ (eBioscience) and c-Kit^+/-^ (eBioscience). To enrich for specific IEC populations, cells were isolated from Lgr5-GFP mice, stained with the antibodies mentioned above and gated for GFP-high (stem cells), GFP-low (TAs), GFP^-^/CD24^+/-^/c-Kit^+/-^ (secretory lineages) or GFP^-^/CD24^-^/EpCAM^+^ (epithelial cells). A population control of 200 cells was sorted into one well and a no-cell control was sorted into another well. After sorting, the plate was sealed tightly with a Microseal F and centrifuged at 800g for 1 min. The plate was immediately frozen on dry ice and kept at -80ºC until ready for the lysate cleanup. Bulk population cells were sorted into an Eppendorf tube containing 100µl solution of TCL with 1% 2-mercaptoethanol and stored at -80ºC.

*Droplet-based scRNA-seq*. Cells were sorted with the same parameters as described for plate-based scRNA-seq, but into an Eppendorf tube containing 50µl of 0.4% BSA-PBS and stored on ice until proceeding to the GemCode Single Cell Platform or the Chromium Single Cell 3’ Library.

##### Plate-based scRNA-seq

*Single cells.* Libraries were prepared using a modified SMART-Seq2 protocol as previously reported [60]. Briefly, RNA lysate cleanup was performed using RNAClean XP beads (Agencourt), followed by reverse transcription with Maxima Reverse Transcriptase (Life Technologies) and whole transcription amplification (WTA) with KAPA HotStart HIFI 2× ReadyMix (Kapa Biosystems) for 21 cycles. WTA products were purified with Ampure XP beads (Beckman Coulter), quantified with Qubit dsDNA HS Assay Kit (ThermoFisher), and assessed with a high sensitivity DNA chip (Agilent). RNA-seq libraries were constructed from purified WTA products using Nextera XT DNA Library Preparation Kit (Illumina). On each plate, the population and no-cell controls were processed using the same method as the single cells. The libraries were sequenced on an Illumina NextSeq 500.

##### Droplet-based scRNA-seq

Single cells were processed through the GemCode Single Cell Platform using the GemCode Gel Bead, Library and Chip Kits, or the Chromium Single Cell 3’ Library, Gel Bead and Chip Kits (10X Genomics, Pleasanton, CA), following the manufacturer’s protocol. Briefly, an input of 6,000 cells was added to each channel of a chip with a recovery rate of 1,500 cells. The cells were then partitioned into Gel Beads in Emulsion (GEMs) in the GemCode instrument, where cell lysis and barcoded reverse transcription of RNA occurred, followed by amplification, shearing and 5’ adaptor and sample index attachment. Libraries were sequenced on an Illumina NextSeq 500.

##### Div-Seq

Lgr5-GFP mice were intraperitoneally (IP) injected with 100 mg kg^-1^ EdU (Click-iT Plus EdU Pacific Blue Flow Cytometry Assay Kit, Thermo Fisher Scientific) for 2 hours and then sacrificed. Crypts were isolated as described above and Lgr5^High^ cells were FACS sorted into PBS, spun down to remove the supernatant, flash frozen and stored in - 80ºC. Nuclei were then isolated using EZ Prep NUC-101 (Sigma) per manufacturer’s recommendation, and then incubated in the Click-iT Cocktail per manufacturer’s recommendations for 30 min, washed in 1% BSA-PBS and counterstained with Vybrant DyeCyle Ruby stain (Thermo Fisher Scientific) for 15 min. Nuclei were then individually sorted into the wells of 96 well plates with TCL+1% 2-mercaptoethanol as described before [61] using FACS, based on positive Ruby and either EdU^high^ or EdU^low^. Plate-based single-nucleus RNA-seq (snRNA-Seq) was then performed as described above for scRNA-seq.

##### Immunofluorescence and single-molecule fluorescence *in situ* hybridization (smFISH)

*Immunofluorescence (IFA) and immunohistochemistry (IHC):* Staining of small intestinal tissue was conducted as previously described [62]. Briefly, tissues were fixed for 14 hours in formalin, embedded in paraffin and cut into 5 μm thick sections. Sections were deparaffinized with standard techniques, incubated with primary antibodies overnight at 4ºC, and then incubated with secondary antibodies at room temperature for 30 min. Slides were mounted with Slowfade Mountant+DAPI (Life Technologies, S36964) and sealed.

*Single-molecule fluorescence in situ hybridization (smFISH):* RNAScope Fluorescent Multiplex and RNAScope Multiplex Fluorescent v2 (Advanced Cell Diagnostics) were used per manufacturer’s recommendations with the following alterations. Target Retrieval boiling time was adjusted to 12 minutes and incubation with Protease IV at 40ºC was adjusted to 8 minutes. Slides were mounted with Slowfade Mountant+DAPI (Life Technologies, S36964) and sealed.

*Combined IFA and smFISH* was implemented by first performing smFISH, as described above, with the following alterations. After Amp 4, tissue sections were washed in washing buffer, incubated with primary antibodies overnight at 4ºC, washed in 1x TBST 3 times and then incubated with secondary antibodies for 30 min at room temperature. Slides were mounted with Slowfade Mountant+DAPI (Life Technologies, S36964) and sealed.

##### Image analysis

*Image acquisition*. Images of tissue sections were taken with a confocal microscope Fluorview FV1200 using Kalman and sequential laser emission to reduce noise and signal overlap. Scale bars were added to each image using the confocal software FV10-ASW 3.1 Viewer. Images were overlaid and visualized using ImageJ software [63].

*Quantification of proliferating stem cells.* Combined IFA and smFISH images of wild-type C57BL/6J small intestinal tissues were assessed by staining for E-Cadherin to mark cell borders, the canonical proliferation marker *mKi67*, and either the common ISC marker *Lgr5*, our predicted lcISC markers (*Cyp2e1* or *Fgfr4*) or our predicted hcISC markers (*Psrc1* or *Cenpf*). A line was drawn to establish the bottom of the crypt, termed “stem cell zone”, and quantification was only assessed within that zone. For each ISC subset marker, more than 10 randomly chosen intact crypts were analyzed. Cells were examined by double blind quantification and were determined double positive if they co-expressed *mKi67* and one of the ISC subset markers. Proliferating cells in each ISC subset was measured by calculating the fraction of double positive cells out of all cells positive for the specific ISC subset marker.

*Automated quantification of Lgr5 mRNA molecules in smFISH images* of intestinal crypts within different mouse models (**Figure 4** and **Figure S9**) was performed using a custom Python script.

##### Antibodies and probes

*Antibodies used for immunofluorescence*. Rabbit anti-DCLK1 (1:200, Abcam GR245168-1), mouse anti-E-cadherin (1:100, BD Biosciences 610181), rat anti-Lysozyme (Dako, A0099) and anti-mouse I-A/I-E (1:100, Biolegend 107601). Alexa Fluor 488-, 594-, and 647-conjugated secondary antibodies were used (Life Technologies).

*Probes used for single-molecule RNAscope* (Advanced Cell Diagnostics): Lgr5 (C1,C3), Cyp2e1 (C2), Psrc1 (C1), Fgfr4 (C2), Cenpf (C3), mKi67 (C1,C3).

##### Th cell polarization *in vitro*

CD4^+^ naïve (CD44^lo^CD62L^+^CD25^-^) T cells were isolated from spleen and lymph nodes of 7–10 weeks old C57BL/6J mice using flow cytometry cell sorting. The purity of isolated T cell populations routinely exceeded 98%. Naïve T cells were stimulated with plate-bound anti-CD3 (145-2C11, 1mg/ml) and anti-CD28 (PV-1, 1mg/ml) and polarizing cytokines (Th1: 4 ng/ml IL-12; Th2: 4 ng/ml IL-4; Th17: 10 ng/ml IL-6, 2 ng/ml TGF-β1; iT_reg_: 5 ng/ml TGF-β1; all cytokines from R&D).

##### Intestinal organoid cultures

*Organoid cultures*. Following crypt isolation from the whole small intestine [64], the single-cell suspension was re-suspended in Matrigel (BD Bioscience) with 1µM Jagged-1 peptide (Ana-Spec). Roughly 300 crypts embedded in 25µl of Matrigel were seeded onto each well of a 24-well plate. Once solidified, the Matrigel was incubated in 600µl culture medium (Advanced DMEM/F12, Invitrogen) with streptomycin/penicillin and glutamatax and supplemented with EGF (100 ng/mL, Peprotech), R-Spondin-1 (600ng/mL, R&D), Noggin (100ng/mL, Prepotech), Y-276432 dihydrochloride monohydrate (10µM, Tochris), N-acetyl-1-cysteine (1µM, Sigma-Aldrich), N2 (1X, Life Technologies), B27 (1X, Life Technologies) and Wnt3A (25ng/mL, R&D Systems). Fresh media was replaced on day 3, and organoids were passaged by dissociation with TrypLE and re-suspended in new Matrigel on day 6 with a 1:3 split ratio.

*T helper cell co-culture experiments*. Organoids were cultured with Th1, Th2, Th17 or iT_regs_. Roughly 10,000 T helper cells were added to each well of 500 organoids and were supplemented either to the medium or suspended in the Matrigel. Treated organoids were dissociated and subjected to droplet based scRNA-seq.

*Cytokine treated organoids*. Organoids were additionally treated with 0.5u/ml IFNγ, 20 ng/ml IL-13, 20 ng/ml IL-17A or 10ng/ml IL-10 in the culture medium for 3 days.

*Re-seeding after cytokine treatment*. 500 organoids/well were treated with cytokines, as in the cytokine treated organoids above, collected after 3 days and then re-seeded at 500 organoids/well in fresh media without cytokines. Each day, images were taken at 2X magnification and quantification of organoids number was performed with the ImageJ software.

##### Two-photon intra-vital microscopy (2P-IVM) of T cells and ISCs

To generate gut-homing T cells visualized by 2P-IVM, a combination of modified protocols [65, 66] was used. CD4^+^ T cells were isolated from spleen, pLN and mLN from β-actin-RFP mice using a MACS CD4 T cell positive-selection kit (Miltenyi clone L3T4) following the manufacturer’s instructions. Plates were pre-treated with 5 μg/mL anti-CD3 (clone 145-2C11) and 1 μg/mL anti-CD28 (clone 37.51) and 1×10^6^ CD4^+^ T cells were added to each well for a final volume of 2.5mL in complete RPMI1640 media supplemented with all-trans Retinoic Acid (100nM, Sigma R2625). The T cells were cultured for 96 hours before replacing half of the volume with fresh media containing 20U/mL of rIL-2 and then cultured for another 48 hours. Before adoptive transfer into Lgr5-GFP hosts, the gut-homing phenotype was validated with flow cytometry for α4β7 and CCR9 expression. 1×10^7^ cells were then transferred into recipient mice for two hours, and treated with 20ug of anti-CD3 (clone 2C11). 2P-IVM was performed 72 hours following transfer. The small intestine was surgically exposed through a laparotomy incision. Anesthetized mice were placed on a custom-built stage with a loop of the intact small intestine fixed to a temperature-controlled metallic support to facilitate exposure of the serosal aspect to a water-immersion 20X objective (0.95 numerical aperture) of an upright microscope (Prairie Technologies). A Mai Tai Ti:sapphire laser (Spectra-Physics) was tuned between 870nm and 900nm for multiphoton excitation and second-harmonic generation. For dynamic analysis of cell interaction in four dimensions, several X/Y sections (512x512) with Z spacing ranging from 2μm to 4μm were acquired every 15-20 seconds with an electronic zoom varying from 1X to 3X. Emitted light and second-harmonic signals were directed through 450/80-nm, 525/50-nm and 630/120-nm band-pass filters and detected with non-descanned detectors. Post-acquisition image analysis, volume-rendering and four-dimensional time-lapse videos were performed using Imaris software (Bitplane scientific software).

#### Computational Methods

##### Pre-processing of plate-based scRNA-seq data

BAM files were converted to merged, de-multiplexed FASTQs using the Illumina provided Bcl2Fastq software package v2.17.1.14. Paired-end reads were mapped to the UCSC mm10 mouse transcriptome using Bowtie [67] with parameters "-q --phred33-quals -n 1 -e 99999999 -l 25 -I 1 -X 2000 -a -m 15 -S -p 6", which allows alignment of sequences with one mismatch. Expression levels of genes were quantified as transcript-per-million (TPM) values calculated by RSEM [68] v1.2.3 in paired-end mode. For each cell, we quantified the number of genes for which at least one read was mapped, and then excluded all cells with either fewer than 3,000 detected genes or a transcriptome-mapping rate of less than 40%.

Selection of variable genes was performed by fitting a generalized linear model to the relationship between the squared coefficient of variation (CV) and the mean expression level in log/log space, and selecting genes that significantly deviated (*p*<0.05) from the fitted curve, as previously described [69].

For re-analysis of published data [70] (**Figure S5F**) normalized transcript counts were downloaded directly from the published data tables. Cell-quality filtering, transcript count normalization, tSNE, and clustering using the RaceID algorithm [70], were all performed using R scripts published online by the authors, using all default settings.

##### Pre-processing of droplet-based scRNA-seq data

De-multiplexing, alignment to the mm10 mouse transcriptome and UMI-collapsing were performed using the Cellranger toolkit (version 1.0.1) provided by 10X Genomics. For each cell, we quantified the number of genes for which at least one read was mapped, and then excluded all cells with fewer than 800 detected genes. Expression values *E_i,j_* for gene *i* in cell *j* were calculated by dividing UMI count values for gene *i* by the sum of the UMI counts in cell *j*, to normalize for differences in coverage, and then multiplying by 10,000 to create TPM-like values, and finally calculating log_2_(TPM+1) values. Batch correction was performed using ComBat [71] as implemented in the R package sva [72], using the default parametric adjustment mode. The output was a corrected expression matrix, which was used as input to further analysis. We identified highly variable genes as described above.

##### Dimensionality reduction by PCA

We restricted the expression matrix to the subsets of variable genes and high-quality cells noted above, and values were centered and scaled before input to PCA, which was implemented using the R function ‘prcomp’ from the ‘stats’ package for the plate-based dataset. For the droplet-based data, we used a randomized approximation to PCA [73], implemented using the ‘rpca’ function from the ‘rsvd’ R package, with the parameter *k* set to 100. This low-rank approximation is several orders of magnitude faster to compute for very wide matrices. After PCA, significant PCs were identified using a permutation test as previously described [74], implemented using the ‘permutationPA’ function from the ‘jackstraw’ R package. Only scores from these significant PCs were used as the input to further analysis.

##### tSNE visualization

For visualization purposes only (and *not* for clustering), dimensionality was further reduced using the Barnes-Hut approximate version of the *t*-distributed stochastic neighbor embedding (tSNE) [75, 76]. This was implemented using the ‘Rtsne’ function from the ‘Rtsne’ R package using 20,000 iterations and a perplexity setting that ranged from 10 to 30 depending on the size of the dataset. Scores from the first *n* PCs were used as the input to tSNE, where *n* was determined for each dataset using the permutation test described above.

##### Removing doublets

In the plate-based dataset, several cells were outliers in terms of library complexity, which could possibly correspond to more than one individual cell per sequencing library, or ‘doublets’. As a precaution, we removed any cells in the top quantile 1% of the distribution of genes detected per cell, as these may correspond to doublets.

##### *k-*NN graph based clustering

To cluster single cells by their expression profiles, we used unsupervised clustering, based on the Infomap graph-clustering algorithm [77], following approaches recently described for single-cell CyTOF data [78] and scRNA-seq [79]. Briefly, we constructed a *k* nearest neighbor (*k*-NN) graph on the data using as the metric for each pair of cells, the distance between the scores of significant PCs. The parameter *k* was chosen in a manner roughly consistent with the size of the dataset (**Table S1**). Specifically, *k* was set to 600, 200 and 50 for the droplet dataset of 23,177, 4,332 and 1,090 cells from combined T cell and cytokines (**Figure S5D**), IL-13-treated and Th1 co-cultured organoids, respectively. For *in vivo* mouse models (**Figure S8B**), *k* was set to 100, 300, 175, and 100 for nude mice, TCRβ KO, Foxp3-DTR and MHCII^Δgut^ respectively. For sub-clustering of stem cell subsets, we used *k=*150 and *k=*40 for the 637 and 123 Lgr5^+^ stem cells from our plate-based and the previously published [70] datasets, respectively. The combined *Salmonella* and *H*. *polygyrus* infection dataset (**Figure S4B-D**) contained 5,122 immune cells and *k* was set to 200. The *k*-NN graph was computed using the function ‘nng’ from the R package ‘cccd’ and was then used as the input to Infomap [77], implemented using the ‘infomap.community’ function from the ‘igraph’ R package.

Detected clusters were annotated by cell types or states using known markers for IEC subtypes [80]. Specifically, for each known epithelial type we selected five canonical marker genes (*e.g.*, *Lgr5*, *Ascl2*, *Slc12a2*, *Axin2* and *Olfm4* for stem cells, or *Lyz1*, *Defa17*, *Defa22*, *Defa24* and *Ang4* for Paneth cells), and scored all clusters for their expression (see below for signature scoring procedure). In all cases, one cluster unambiguously expressed each cell-type signature, with two exceptions: in the plate-based dataset, two clusters both expressed high levels of ISC markers (**Figure S1A**) and accordingly were merged to form a ‘Stem’ cluster and two other clusters were merged to form a ‘TA’ cluster based on high expression of cell-cycle genes and low-to-moderate expression of ISC genes.

##### Assigning the three ISC states to region of origin using supervised classification

To study the anatomical distribution of ISCs in different parts of the small intestine, we used a classification approach. First, we developed a classifier for the anatomical origin of ISCs using single-cell expression profiles of 2,965 ISCs extracted from duodenum, jejunum and ileum [81], by compiling a discriminative feature set using the expression levels of all genes differentially expressed (FDR < 0.1, Mann-Whitney U-test, log_2_ fold-change > 0.25) between stem cells from the three regions, and also the scores along the first 25 PCs. A ‘random forest’ classifier was trained on these features, and subsequently distinguished between ISCs from the three regions with an average out-of-bag accuracy of 92.9%. Finally, we used the trained classifier to classify the 637 ISCs (**Figure 1**) and infer the fraction of cells drawn from each intestinal region found in each ISC state (**Figure S1F**).

##### Cell-cell similarity matrix

To visualize heterogeneity of ISCs within the ‘Stem’ cluster (637 cells), cell-cell similarities were computed. Principal component (PC) scores for each cell were computed across the 637 cells using the R function ‘prcomp’ as described above. The distance between cell *i* and *j* was calculated as the Pearson correlation between the scores of these two cells along the first 10 PCs. This distance matrix was then hierarchically clustered using Ward’s method, implemented using the R function ‘hclust’ (with the ‘method’ argument set to ‘ward.D2’), and visualized as a heatmap using the R function ‘aheatmap’ (**Figure 1B**).

##### Cell-cycle and ISC subset signatures

To identify maximally specific genes associated with the three ISC subsets, we performed differential expression tests between each possible pairwise comparison between clusters. To ensure specificity of the detected marker genes to stem cells, the set of clusters included both the three ISC subsets (3 clusters), and all other detected IEC clusters (8 clusters; **Figure S1A**, right) for a total of 11 clusters. Then, for a given cluster, putative signature genes were filtered using the maximum FDR Q-value and ranked by the minimum log_2_(fold-change). The minimum fold-change and maximum Q-value represents the weakest effect-size across all pairwise comparisons, therefore this is a stringent criterion. ISC subset signatures (**Table S2**) were obtained using a maximum FDR of 0.25 and a minimum log_2_(fold-change) of 0.25. To exclude the explicit effect of known cell-cycle genes on the gene signature of the ISC subsets we filtered out any gene annotated as directly participating in cell-cycle regulation. Annotated cell-cycle genes were downloaded from the gene ontology (GO): http://amigo.geneontology.org/amigo/term/GO:0007049, and any gene appearing on this list was removed from the signature gene sets.

Gene sets associated with G1/S and G2/M phases of the cell-cycle were downloaded from http://www.cell.com/cms/attachment/2051395126/2059328514/mmc2.xlsx [82]. A set of cell-cycle genes to assess overall proliferation (see below for scoring procedure) was defined as the union of the G1/S and G2/M sets.

##### Scoring cells using signature gene sets

To score a specific set of *n* genes in a given cell, a ‘background’ gene set was defined to control for differences in sequencing coverage and library complexity between cells [83]. The background gene set was selected to be similar to the genes of interest in terms of expression level. Specifically, the 10*n* nearest gene neighbors in the 2-D space defined by mean expression and detection frequency across all cells were selected. The signature score for that cell was then defined as the mean expression of the *n* signature genes in that cell, minus the mean expression of the 10*n* background genes in that cell.

##### Testing for shifts in cell proportions in intestinal organoids

Under several conditions, we observed dramatic changes in the frequency of epithelial cell subtypes (**Figure 3**). The statistical significance of these shifts was assessed by calculating, for each condition comparison and cell type, the exact hypergeometric probability (without replacement) of the observed change in cell numbers.

Specifically, given that ***m*** and ***n*** total cells (of all cell types) are sequenced in a treatment and control condition respectively, we test, for a given cell type, whether the number of ***k*** and ***q*** of observed cells of type ***C*** in total and treatment condition respectively, significantly deviates from a null model given by the hypergeometric distribution. The probability of observing these values was calculated using the R function ‘phyper’ from the ‘stats’ package, using the command:
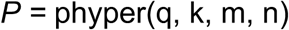
and was reported as a hypergeometric *p-*value.

##### Testing for shifts in cell proportions *in vivo*

In the case of *in vivo* perturbation experiments (**Figure 4**), we used a regression model to control for any mouse-to-mouse variability amongst our biological replicates. For each cell-type, we model the number of cells detected in each analyzed mouse as a random count variable using a negative binomial distribution. The *rate* of detection is then modeled by using the natural log of the total number of cells profiled in a given mouse as an offset variable. The condition of each mouse (*i.e.*, knock-out or wild-type) was provided as a covariate. The model was fit using the R command ‘glm’ from the ‘stats’ package. The *p*-value for the significance of the effect produced by the knock-out was then assessed using a Wald test.

##### GO analysis

GO analysis was performed using the ‘goseq’ R package [84], using significantly differentially expressed genes (FDR <0.05) as target genes, and all genes expressed with log_2_(TPM+1) > 3 in at least 10 cells as background.

### Supplementary Figure legends

**Figure S1.**
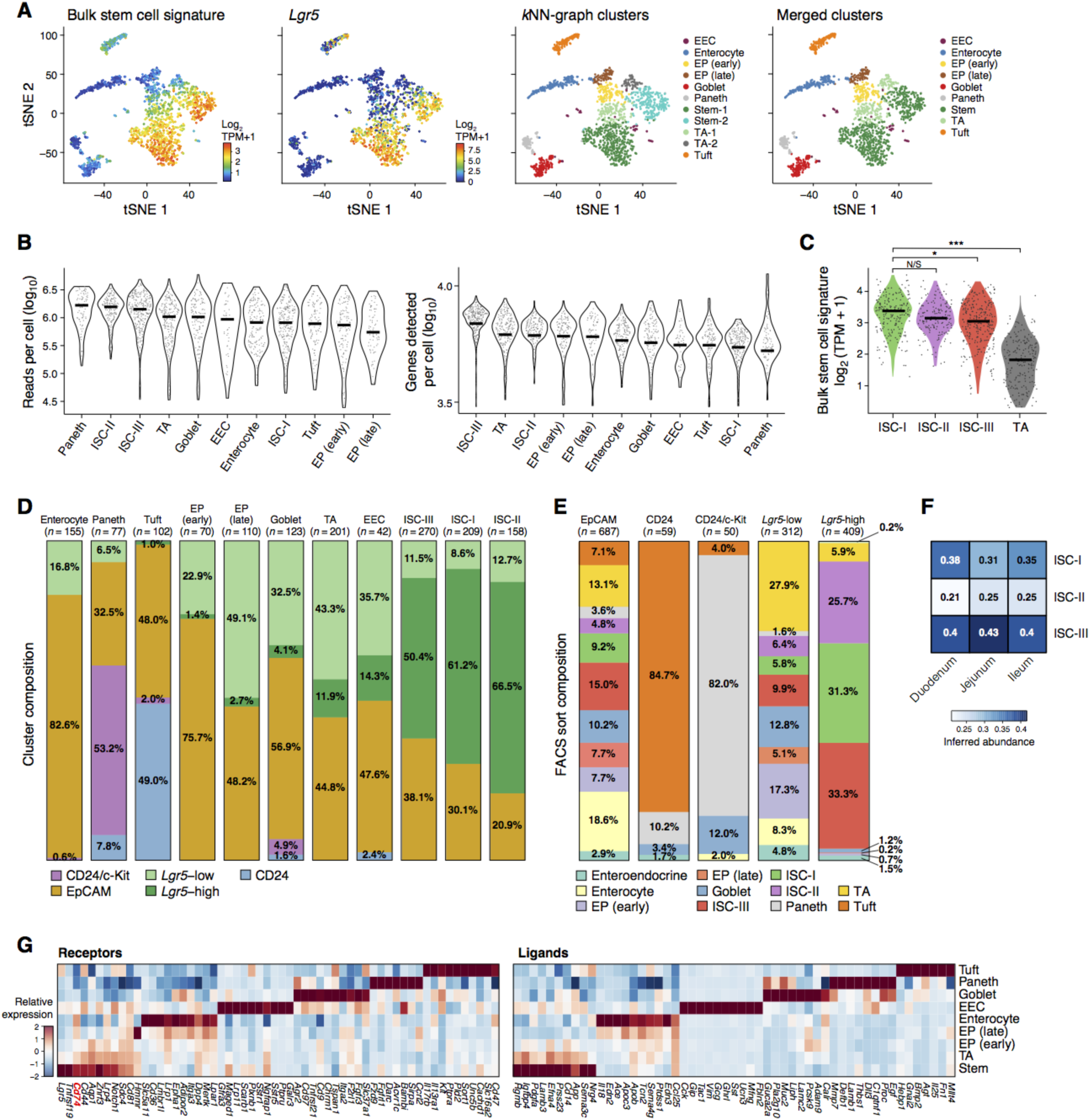
Identification of *Lgr5*^+^ stem cells by single cell RNA-seq. **A.** Intestinal stem cells (ISCs) identified from scRNA-seq data by unsupervised clustering and *post-hoc* annotation. tSNE visualization of 1,522 single cells (points) profiled by full-length scRNA-seq [81]. Cells are colored by the mean expression (mean Log_2_(TPM+1), color bar) of a previously published [85] ISC gene signature (left), the marker gene *Lgr5* (log_2_(TPM+1), color bar, center left), or by a color code of clusters from *k*NN-graph clustering (center right), which identifies two clusters (Stem-1 and Stem-2, dark green and cyan, respectively) both of which are positive for the ISC signature genes and express *Lgr5*. The union of these two clusters (dark green, right) forms a set of 637 ISCs, which were used for further analyses. EP: Enterocyte progenitor, EEC: enteroendocrine cell. **B.** Quality control. Violin plots show the distributions of reads per cell (*y*-axis, left) and genes detected per cell (*y*-axis, right) in each IEC-type (*x*-axis, as defined in (**A**)). Horizontal bars: median. **C.** All ISC subsets express a stemness signature. Violin plot showing mean expression (log_2_(TPM+1), *y*-axis) of stem cell signature genes [85], in each of the three ISC subsets as well as in the cluster of 201 TA progenitors (**Figure S1A**) (*x*-axis). * *p*<0.001, ** *p*<1x10^-5^, *** *p*<1x10^-7^ (Mann-Whitney U-test). **D,E.** Validation of *in silico* cell type identification using FACS. (**D**) Proportion (percentage, y-axis) of cells from FACS sorted EpCAM^*+*^, Lgr5^Low^, Lgr5^High^, CD24^+^ and CD24^+^/c-Kit^+^ fractions (color legend) in each of the cell type clusters (bars), identified in the 1,522 IECs sequenced using full-length scRNA-seq (as in (**A)**, **Methods**). At least 50% of the cells in each of the ISC subsets are Lgr5^High^, while less than 15% of the cells in any other subset are Lgr5^High^. (**E**) Proportion (percentage, *y*-axis) of cells from each identified cluster (color legend) in each of the FACS fractions (bars). 90.3% of cells in the Lgr5^High^ fraction are assigned to one of the three ISC states. **F.** Three ISC subsets are similarly represented along the small intestine. Heatmap shows the fraction (color legend) of cells in each of the detected ISC states (rows) among the ISC isolated from each of three spatial regions (columns), as inferred using a random forest classifier trained on 2,965 ISCs [81], extracted from each of the respective gut regions (**Methods**). **G.** Cell type-enriched ligands and receptors. Average relative expression (Z-score of mean log_2_(TPM+1), color bar) of the top 10 receptors (left) and ligands (right, columns) enriched (FDR<0.05, Mann-Whitney U-test) in each cell type (rows). The invariant chain of MHCII, *Cd74*, is highlighted in red (left).

**Figure S2.**
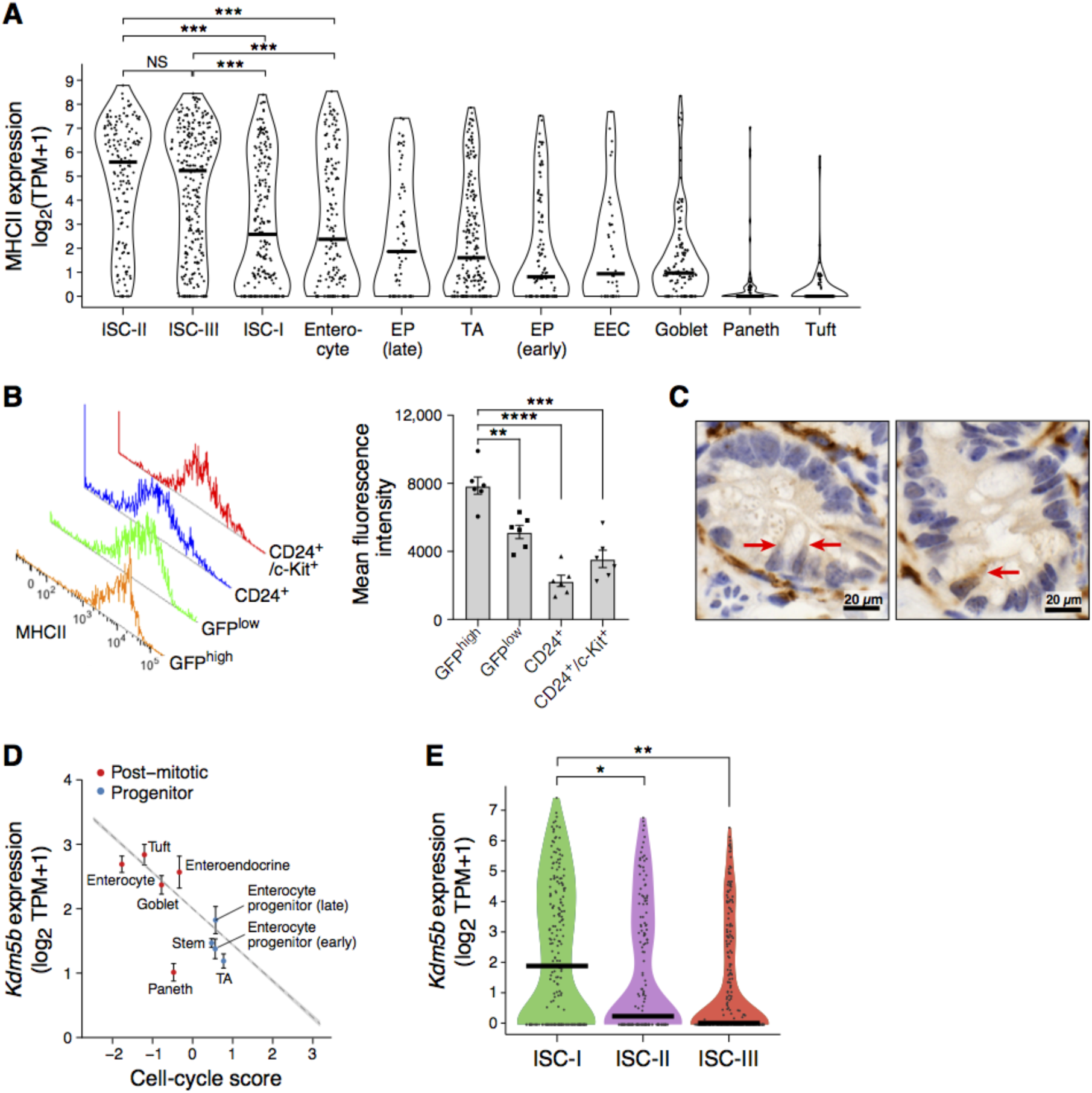
Identification and characterization of MHCII-expressing *Lgr5*^+^ stem cells. **A.** MHCII signature is largely restricted to ISCs. Violin plot shows the distribution of mean expression levels (log_2_(TPM+1), *y*-axis, bar: median) of MHCII genes (*H2-Ab1, H2-Aa, Ciita, Cd74, H2-DMa, H2-DMb1*) in IEC types (**Figure S1A**) from the 1,522 IECs profiled by full-length scRNA-seq. EP: Enterocyte progenitor, EEC: enteroendocrine cell. **B.** Protein-level quantification via FACS of MHCII expression in IECs from Lgr5-GFP mouse. Distribution (left) and mean (bar plot, right) of detected fluorescence corresponding to MHCII protein expression in populations sorted for GFP-high, GFP-low, CD24^+^ or CD24^+^/c-kit^+^. Error bars: SEM. *n*=6 mice. **C.** MHCII is expressed in intestinal crypts of wild type mice. IHC images show MHCII expression (I-A/I-E, brown) within crypts of WT mice (*n*=2 mice). Red arrow, MHCII^+^ cell. Scale bar, 20µm. **D,E.** Known quiescence marker *Kdm5b* identifies post-mitotic cells and low-cycling ISC-I subset. (**D**) Scatter plot shows the negative relationship between cell-cycle score (*x-*axis) and the mean expression of the putative quiescence marker *Kdm5b* [86-90] (log_2_(TPM+1), y-axis) in each IEC-type (dots) for both proliferating (blue) and post-mitotic (red) cells. Trend line shows the random-effects linear model fit to all 1,522 cells. Error bars: SEM. (**E**) Violin plot of the distribution of expression level of *Kdm5b* (log_2_(TPM+1), *y*-axis) in ISC-I, ISC-II, and ISC-III clusters (*x* axis) (* *p*<0.05, ** *p*<0.005, Mann-Whitney U test).

**Figure S3.**
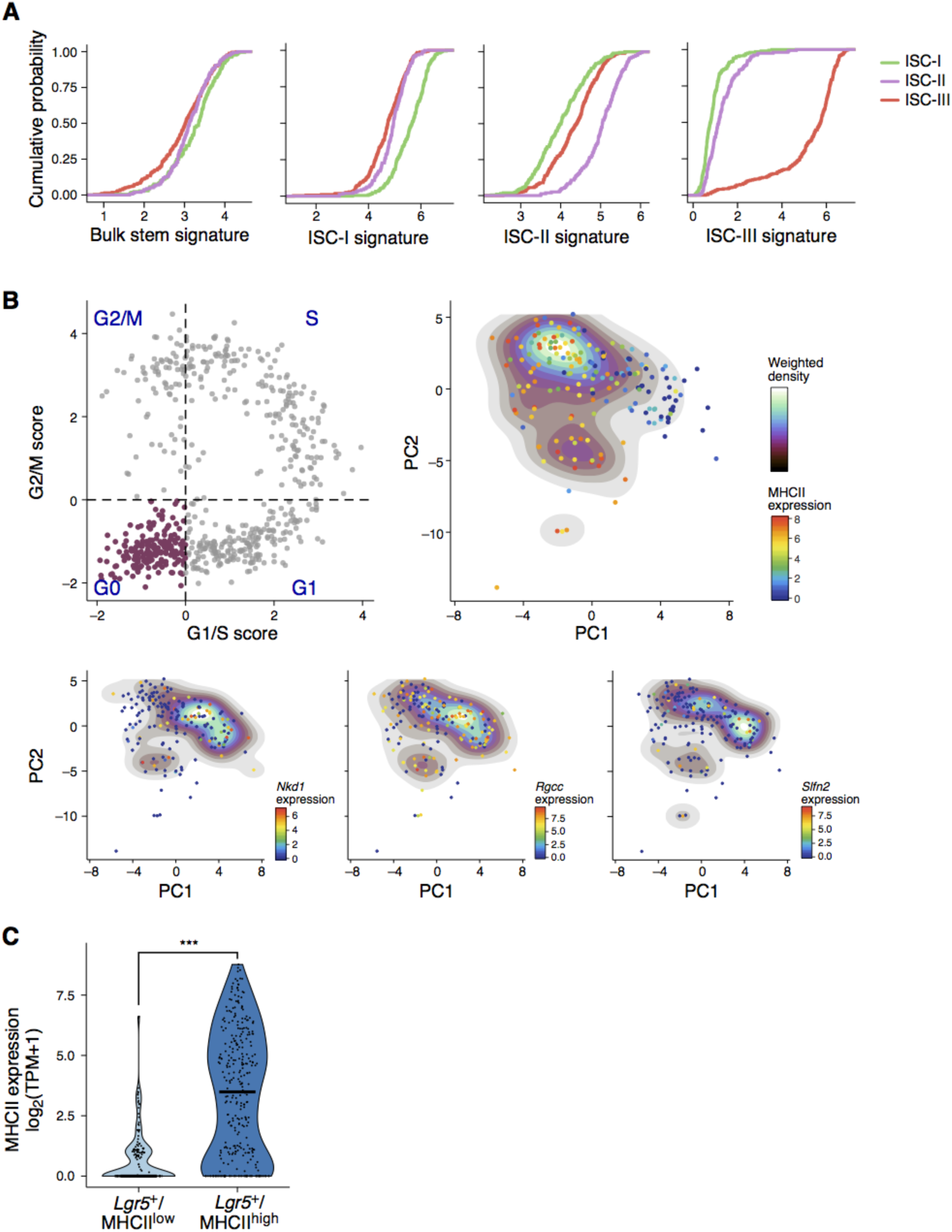
ISC states are distinguishable beyond proliferation. **A.** Distinct signatures of the three ISC subsets. Shown are cumulative distribution functions (CDF) of signature scores (*x*-axis) in the cells from each of the three subsets of ISCs (colored curves) of a published ISC signature from bulk data [85] (left), or signatures of 25 genes defined by differential expression testing in each subset (**Methods**). **B**. Two ISC subsets found in G0 are varying by MHCII expression. Top left: Scatter plot of the G1/S (*x*-axis) and G2/M (*y*-axis) signature scores for 637 Lgr5^+^ ISCs (points). A subset of 183 Lgr5^+^ ISCs that are likely in G0 is marked in purple. Other panels: PCA of these 183 ISCs, where cells (dots; density marked by contours) are colored by the expression (log_2_(TPM+1), color bar) of MHCII (top right panel) or by three ISC-I markers (bottom panels). Two subsets of cells are separated by PC-1: one is MHCII^low^ and positive for ISC-I markers and the other is MHCII^high^. **C.** MHCII^high^ ISCs sorted by FACS express higher levels of MHCII mRNA. Violin plot shows the distribution of the mean expression level (*y*-axis, (log_2_(TPM+1)) of the MHCII gene signature (**Methods**) in each cell (dot) in Lgr5^high^ cells sorted on MHCII^low^ and MHCII^high^ (*x*-axis) (*** *p*<0.0005, Mann-Whitney U test).

**Figure S4.**
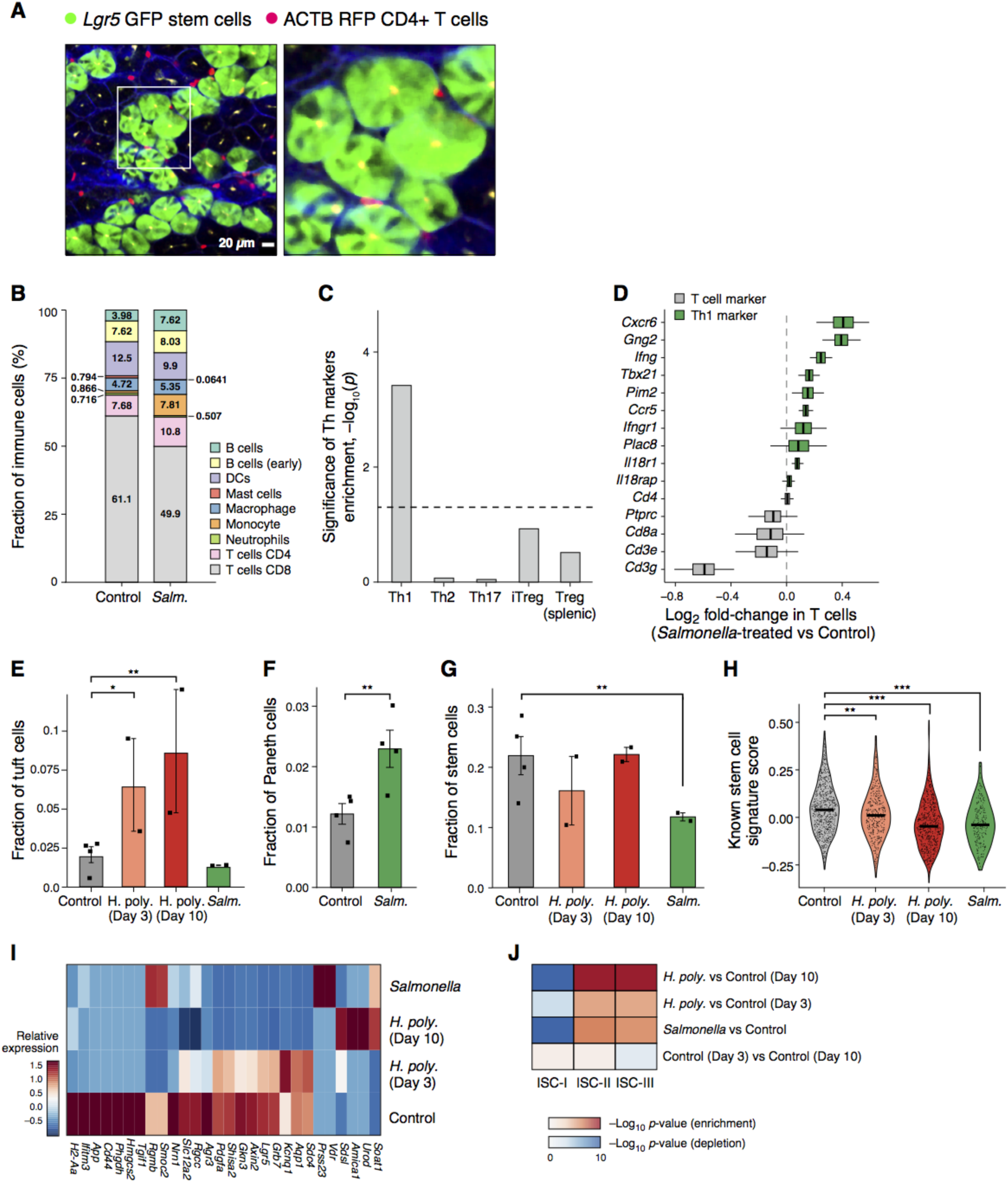
Changes in immune cells, IEC cell-type composition and ISC gene expression in response to pathogen infection *in vivo*. **A.** CD4^+^ T cells interact with stem cells *in vivo*. Two-photon microscopy image of the small intestine from *Lgr5*-GFP (green) knock-in mouse engrafted with RFP^+^ CD4^+^ T cells (red). CD4^+^ T cells are visible in close proximity to Lgr5^+^ ISCs (inset and right). Scale bar, 20µm. **B-D.** *Salmonella enterica* infection induces Th1 polarization in the gut. (**B**) Changes in immune cell proportions. Stacked bar plots show the percentage (*y*-axis) of different immune cell subsets (color legend), as determined by scRNA-seq of 5,122 CD45^+^ cells from the lamina propria of control and *Salmonella*-infected mice. (**C**) Bar plot shows the significance of the enrichment (-log_10_(*p*-value), *y*-axis, hypergeometric test) of marker genes for different T helper subsets (*x*-axis) among the genes induced (FDR<0.05, likelihood-ratio test) in T cells from *Salmonella* infected *vs*. control mice. Dashed line: *p*=0.05. (**D**) Plot shows differential expression (*x*-axis) for each gene (*y*-axis) across 824 T cells from *Salmonella*-infected mice (*n*=4) and 543 T cells from control mice (*n*=5). Bar indicates Bayesian bootstrap [91] estimates of log_2_(fold-change), and hinges and whiskers indicate 25% and 95% confidence intervals, respectively. Th1 cell markers are labeled in green. Dashed line: no differential expression. **E,F.** Changes in fractions of tuft and Paneth cells within the intestinal epithelium after infection. Bar plots show the frequencies of tuft (**E**) and Paneth (**F**) cells (*y*-axis), as determined by unsupervised clustering of droplet-based scRNA-seq data in mice under different conditions (*x*-axis) [81], *n*=2 and 4 mice (points) per group, in (**E**) and (**F**), respectively. * FDR < 10^-5^; ** FDR < 10^-10^, likelihood-ratio test, **Methods**. **G.** Fraction of ISCs within the intestinal epithelium after infection. Cell-type frequencies (*y*-axis) determined by unsupervised clustering of droplet-based scRNA-seq data in each infection model and control mice (*x*-axis, *n*=2 mice (points) per infection group, *n*=4 for control; ** FDR<10^-10^ likelihood-ratio test, **Methods**). **H.** Reduced stemness scores in ISCs during pathogenic infection *in vivo.* Violin plot shows the distribution of the bulk stemness signature score (*y*-axis, **Methods**) of 1,857 ISCs identified by clustering (**Methods**) of 9,842 cells sequenced using droplet-based scRNA-seq from *Salmonella enterica*- or *H. polygyrus*-treated mice and controls [81] (*x*-axis). * *p*<0.01, ** *p<*10^-5^ (Mann-Whitney U-test). **I.** Pathogenic infection reduces the expression of ISC marker genes. Heatmap shows the mean expression (column-wise Z-score of mean log_2_(TPM+1) values, color bar) of all of the known ISC marker genes [85] (columns) that are differentially expressed (FDR<0.05) by 1,857 ISCs, as determined by unsupervised clustering from a total of 9,842 cells profiled by droplet-based scRNA-seq, in control and pathogen-infected mice (rows). **J.** Shifts in the three ISC subsets under infection *in vivo*. Heatmap shows the significance of changes of expression scores (log_10_(*p*-value), Mann-Whitney U-test, color bar, of enrichment (red) and depletion (blue)) within 1,857 ISCs. ISCs in each condition were scored for expression of each ISC subset gene signature (columns, sets and signatures as in **Figure 2**), and the distribution of scores was compared to that in ISCs from control mice (top three rows). Bottom row: comparison between control mice on day 3 and 10 of the *H. polygyrus* infection course.

**Figure S5.**
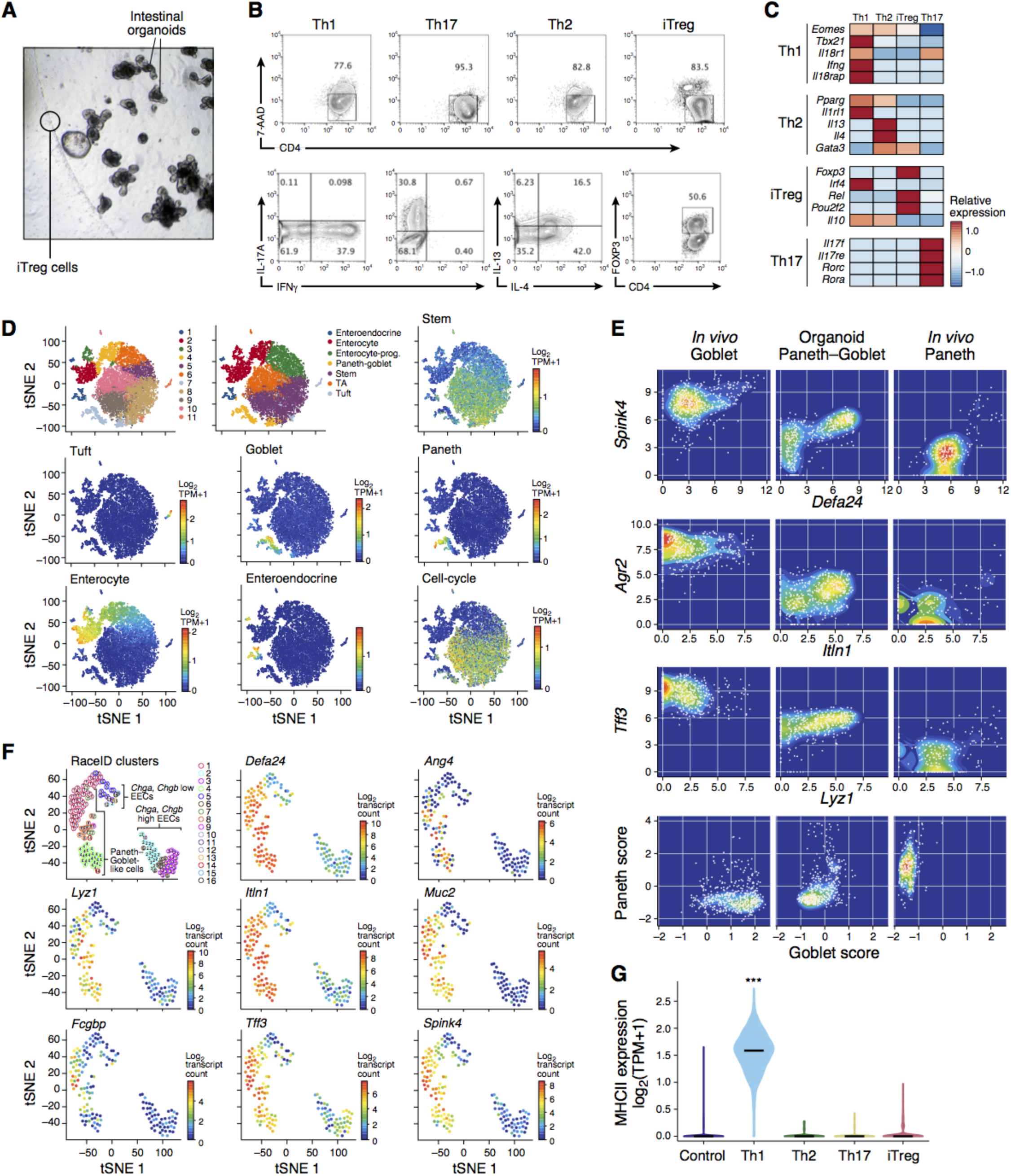
Characterization of intestinal organoids co-cultured with T helper (Th) cells or treated with their key cytokines. **A.** Intestinal organoid co-cultured with T cells. Images of organoid and T_reg_ co-cultures at ×4 magnification. Lines mark T_regs_ and organoids. **B,C.** Validation of *in vitro*-polarized Th cell populations. FACS plots for each of the four subsets of Th cells. Top panels: quantification of the viability dye 7-AAD (*y*-axis) and the Th cell marker CD4 (*x*-axis). Bottom panels: quantification of relevant marker proteins for each Th subset (*x*- and *y*-axes, bottom panels). (**C**). Heatmap shows the mean relative expression (row-wise Z-score of mean log_2_(TPM+1) values, color bar) of canonical marker genes (rows) in the cells from each *in vitro*-differentiated Th cell subset (columns), identified by droplet-based scRNA-seq of co-cultures (**Methods**). **D.** IEC type identification within intestinal organoid cultures. tSNE embeddings of 17,755 single IECs (individual points) isolated from control, cytokine-treated and Th cell co-cultured intestinal organoids and sequenced using droplet-based scRNA-seq. Most cells were merged into a single dataset to maximize the statistical power of clustering (**Methods**). Top left panel: cells colored by cluster assignment from unsupervised *k*NN-graph clustering (**Methods**). Top middle panel: cells colored by *post-hoc* annotations using cell-type signatures derived from *in vivo* scRNA-seq data (clusters that expressed high levels of the same signatures were merged to a final set of seven clusters). All other panels: cells colored by the mean expression (color bar, log_2_(TPM+1)) of the noted cell-type specific signatures. **E.** Organoid-derived secretory IECs co-express markers for goblet and Paneth cells. Scatter plots show the expression levels (top three rows, log_2_(TPM+1)) of canonical markers for goblet cells (*Spink4, Agr2, Tff3, y*-axis) and for Paneth cells (*Defa24, Itln1, Lyz1, x*-axis) or signature scores for goblet and Paneth cells (bottom row, 50 genes), for *in vivo* goblet cells (left), cells in the Paneth-goblet cluster from control organoids (middle) and *in vivo* Paneth cells (right). **F.** Reanalysis of published scRNA-seq data confirms Paneth and goblet cell marker co-expression in organoids. tSNE embeddings of 161 Reg4^+^ cells sorted from intestinal organoid cultures in an independent study [70]. Top left: Cell types were identified using the RaceID clustering algorithm, as in the original publication [70] (colored and numbered nodes). Remaining plots: cells are colored by the expression (log_2_(normalized transcript count), color bars) of canonical markers of Paneth and goblet cells. A group of Paneth-goblet-like cells is clearly observed, where individual cells are double-positive for markers of both cell types. **G.** Induction of MHCII expression in organoids co-cultured with Th1 cells. Violin plot shows the distribution of mean expression levels (log_2_(TPM+1), *y*-axis, bar denotes the median value) of six MHCII genes (*H2-Ab1, H2-Aa, Ciita, Cd74, H2-DMa, H2-DMb1*) in IECs profiled by droplet-based scRNA-seq from control organoids and those co-cultured with each subset of Th cells (total of 6,234 cells, x-axis). *** *p* < 10^-10^, (Mann-Whitney U-test).

**Figure S6.**
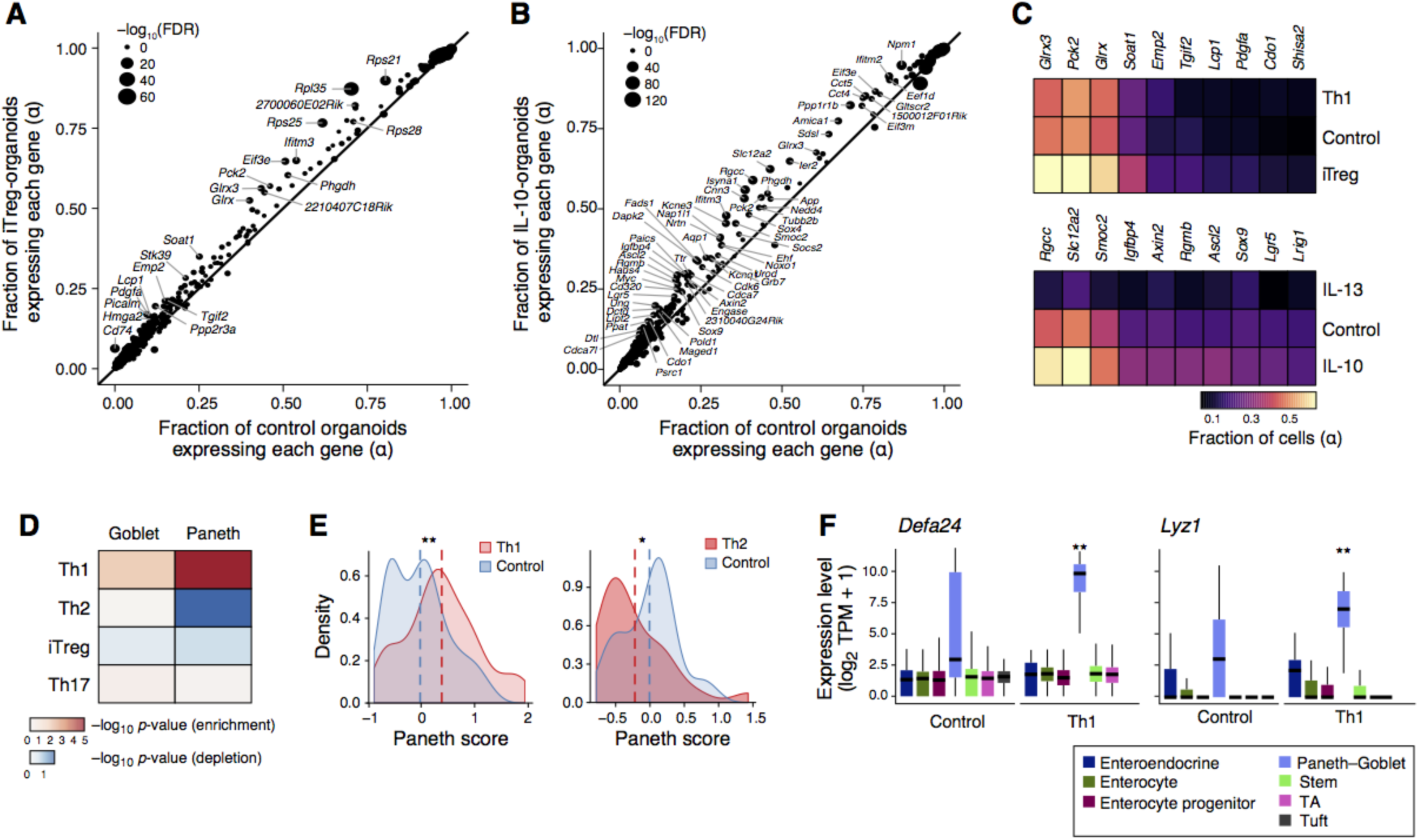
Changes in cell proportions and ISC expression programs in organoids co-cultured with Th subsets or their signature cytokines. **A-C.** Changes in proportion of cells expressing stem cell marker genes after co-culture with iT_regs_ or treatment with IL-10. **A,B.** Scatter plots compare the fraction of cells showing non-zero expression (*α*, *y*-axis) of each gene (dot) in organoids (**A**) co-cultured with iT_reg_-cells or (**B**) treated with IL-10, compared to the fraction in matching control organoids (*α*, *x*-axis). All genes up-regulated (FDR<0.05) are shown, sized relative to their significance (-log_10_(FDR), legend top left). Several key ISC marker genes (**Methods**) are labeled. The diagonal line indicates no change relative to the control organoid. (**C**) Heatmap shows the fraction of cells with non-zero expression (*α*, color bar) of 10 selected stem cell marker genes (columns) within organoids co-cultured with T_regs_ or Th1 cells (top, rows) or treated with IL-13 or IL-10 cytokines (bottom, rows). All genes are expressed at a larger proportion of cells (FDR < 0.05, Mann-Whitney U-test) in organoids co-cultured with iT_reg_ cells (top) or treated with IL-10 (bottom), compared to control organoids. **D-F.** Th1 co-culture up-regulates Paneth cell-related gene expression in organoids. (**D**) Heatmap of the significance of change of the Paneth cell signature score in ‘Paneth-goblet’ cells between different Th co-cultures (rows) and control organoids (-log_10_(*p*-value), Mann-Whitney U-test, color bar, of enrichment (red) and depletion (blue). (**E**) Density histograms of the distribution of Paneth cell signature scores in ‘Paneth-goblet’ cells in organoids co-cultured with either Th1 (red, left) or Th2 (red, right) cells compared to their matching control organoids (blue). Dashed lines denote the mean score. * *p*<0.05, ** *p*<10^-5^ (Mann-Whitney U-test). (**F**) Box plots of the distribution of expression levels (log_2_(TPM+1), *y*-axis) of canonical Paneth cell markers *Defa24* (left) and *Lyz1* (right) in the IEC-type cluster (*x*-axis) from organoids co-cultured with Th1 cells (right part of each panel) compared to control organoids (left part). ** *p*<10^-5^ (Mann-Whitney U-test).

**Figure S7.**
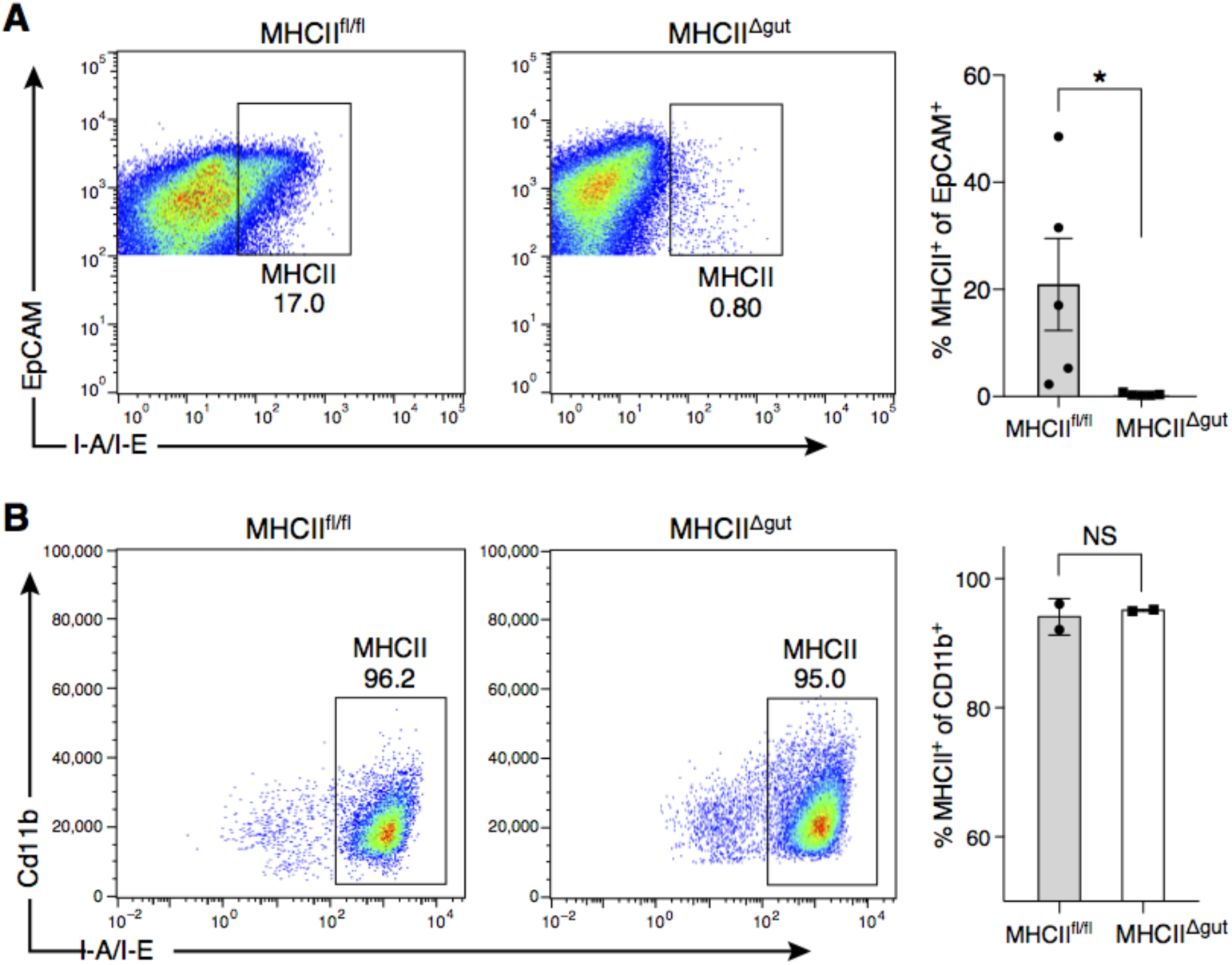
Validation of MHCII knockout in intestinal epithelial cells. **A,B.** Validation of epithelial-specific MHCII knockout by FACS quantification of MHCII-expressing IECs in small intestine (**A**) and mesenteric lymph node **(B)**. Scatter plots (left) and bar plots (right) show the fraction of EpCAM^+^/MHCII^+^ (**A**) or CD11b^+^/MHCII^+^ (**B**) cells in MHCII^fl/fl^ and MHCII knockout mice (MHCII^Δgut^). (**A**) *n*=5 mice, * *p*<0.05. (**B**) *n*=2 mice, NS: not statistically significant.

**Figure S8.**
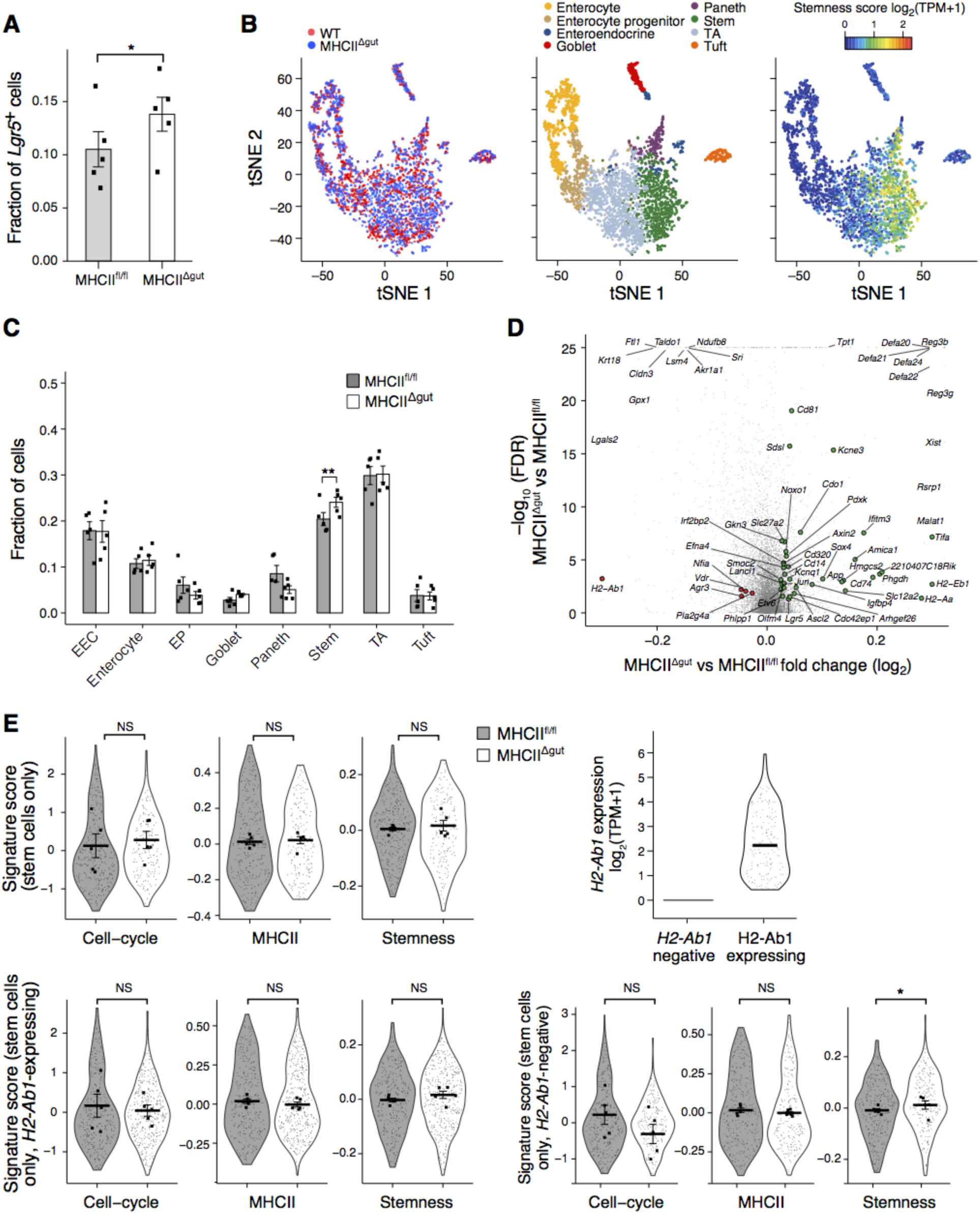
Impact of MHCII knockout in gut epithelial cells on the ISC pool. **A.** Increased proportion of *Lgr5*^+^ cells in MHCII^Δgut^ mice. Bar plot shows the fraction of cells (*y*-axis) in which the transcript for *Lgr5* is detected, amongst the 1,559 cells profiled from MHCII^Δgut^ mice (*n*=5) and 1,617 cells profiled from matched MHCII^fl/fl^ controls (*n*=5). Error bars: SEM, * *p*<0.05, likelihood-ratio test. **B.** scRNA-seq of IECs from MHCII^Δgut^ and matched controls. tSNE embedding of 3,176 cells colored by their genotype (color legend, left), assignment to cell types by unsupervised clustering (middle, **Methods**) and stemness score (right; score as in **Figure S1A**). **C,D.** Expansion of ISCs following KO of MHCII specifically in gut epithelial cells. Bar plot (**C**) and volcano plot (**D**) based on all 1,559 cells in MHCII^Δgut^ mice (*n*=5) *vs*. 1,617 cells from matched MHCII^fl/fl^ controls (*n*=5). Green dots in (**D**): up-regulated ISC genes, red dots: down-regulated ISC genes (FDR<0.05, likelihood-ratio test), grey dots: non-DE genes. **E.** Higher stemness signatures in ISCs from MHCII^Δgut^ mice. Violin plots of the distribution of the signature scores for the cell-cycle (as in **Figure 2A**), MHCII genes and stemness [85] (as in **Figure 1C**), in ISCs from MHCII^fl/fl^ mice (*n*=5, grey) and from MHCII^Δgut^ mice (*n*=5, white), when including either all 381 ISCs (top left), only the 173 ISCs that still have detectable mRNA for *H2-Ab1* (bottom left), or only the 208 ISCs that are a confirmed KO: do not have detectable mRNA for *H2-Ab1* (bottom right). Expression of *H2-Ab1* mRNA is also shown in these two groups (top right). Small dots: individual cells; squares: mean per mouse; * *p* < 0.05, likelihood-ratio test.

**Figure S9.**
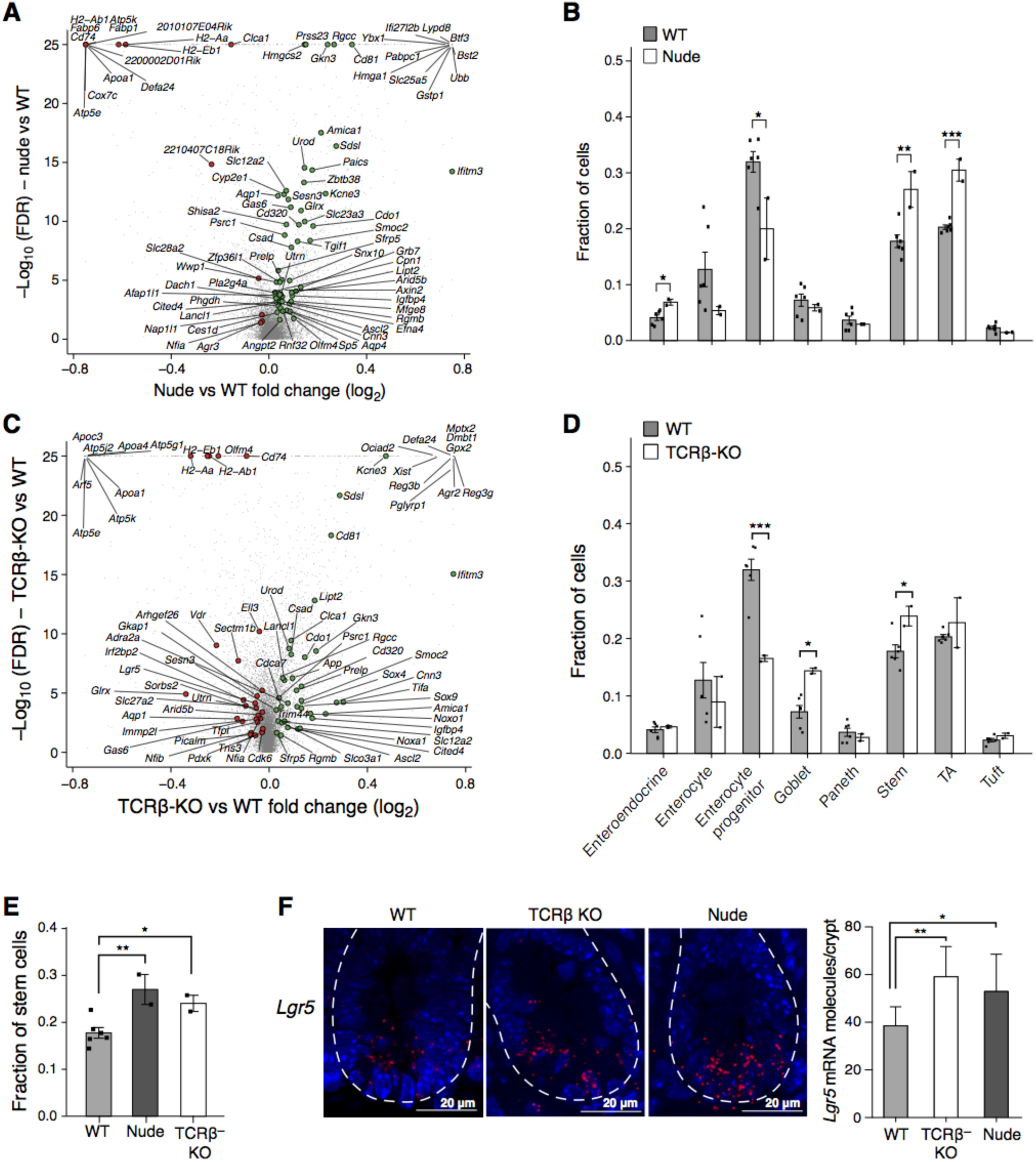
An expanded ISC pool in T cell-depleted mouse models. Volcano plots (**A,C**) show mean log_2_ fold-change (*x*-axis) and significance (-log_10_(FDR), **Methods**) of differential expression between 7,216 cells from WT mice (*n*=6), 2,967 cells from nude mice (**A**, *n*=2 mice) or 9,488 cells from TCRβ-KO mice (**C**, *n*=2 mice). Green dots: up-regulated ISC genes, red dots: down-regulated ISC genes (FDR<0.05, likelihood-ratio test), grey dots: non-DE genes. Bar plots (**B,D**) show frequency (*y*-axis) of each major IEC-type (*x*-axis), as determined by unsupervised clustering (**Methods**), in cells from WT (grey) *vs*. nude (white, **B**) or TCRβ-KO mice (white, **D**). Dots correspond to individual mice. Error bars are SEM. (* FDR<0.05, ** FDR<0.005, *** FDR<10^-5^, likelihood-ratio test). **E,F.** ISC expansion in two T cell-depleted mouse models. (**E**) Bar plot shows the fraction of epithelial cells which are stem cells (*y*-axis), as determined by unsupervised clustering of scRNA-seq profiles, from WT (7,216 cells, *n=*6), nude (2,967 cells, *n=*2), and TCRβ-KO mice (9,488 cells, *n=*2). Dots correspond to individual mice. Error bars are SEM. (* *p*<0.05, ** *p*<10^-3^, *** *p*<10^-5^, likelihood-ratio test, **Methods**). (**F**) smFISH of *Lgr5*^+^ cells in the crypt. Left: *Lgr5* expression (red) in intestinal crypts of wild type (left), TCRβ-KO (center), and nude (right) mice. Right: bar plot shows the number of *Lgr5* molecules detected per crypt (*y-axis*) in each of the three models (*x-axis*). *n=*2 mice and 8 fields per group. Error bars: SD (* *p*<0.05, ** *p*<0.005, *t*-test, **Methods**).

**Figure S10.**
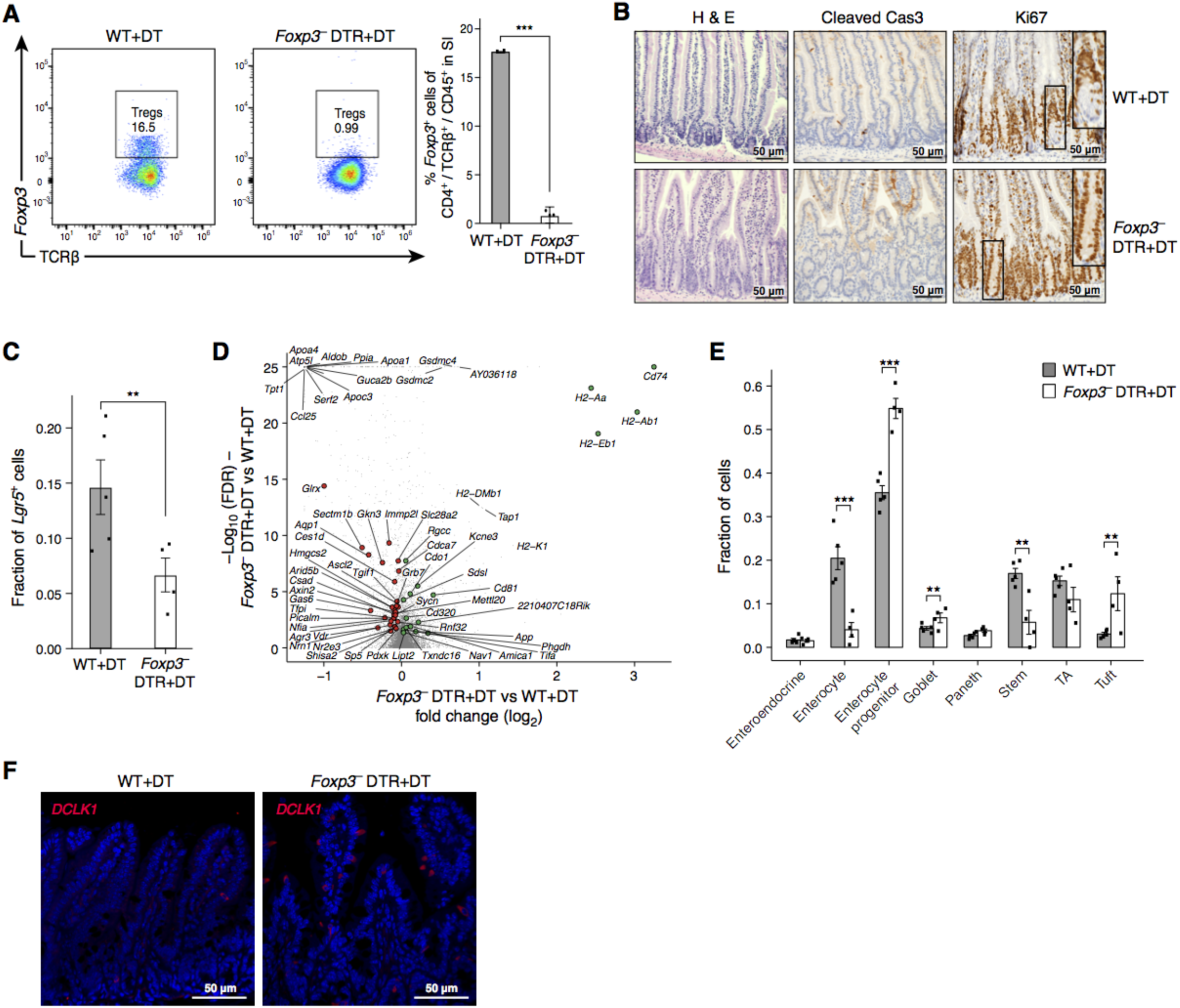
Impact of T_reg_ depletion on ISC pool *in vivo*. **A-E.** Reduction of the ISC pool following T_reg_ depletion in *Foxp3*-DTR mice. **A.** Representative FACS plot (left) and quantified mean proportion (bar plot, right) of TCRβ^+^Foxp3^+^ T_regs_ out of all CD4^+^ TCRβ^+^ cells in the small intestine of WT and *Foxp3*-DTR mice (*y*-axis) after 7 days of DT treatment. Dots: individual mice. Error bars: SEM (*n*=3 mice, *** p<0.0005 *t*-test). **B.** IHC images of FFPE sections stained for H&E (left), cleaved Caspase-3 (brown, middle) and Ki67 (brown, right) in WT (top row) and *Foxp3*-DTR mice (bottom row) after 7 days of DT treatment. Inset is 3× magnification showing T_reg_ depletion results in proliferation at the bottom of the crypts, where stem cells reside, with no signs of apoptosis. Scale bar, 50µm. **C.** Bar plot of the frequency of cells in which *Lgr5* mRNA is detected (*y*-axis) in WT and *Foxp3*-DTR mice both treated with DT. (Error bars: SEM, ** *p*<0.005 likelihood-ratio test). **D.** Volcano plot shows mean log_2_ (fold-change, *x*-axis) and significance (-log_10_(FDR), **Methods**) of differential expression between 815 cells from *Foxp3*-DTR mice (*n*=4) *vs*. 2,572 cells from matched WT controls (*n*=5) both treated with DT. Green dots: upregulated ISC genes, Red dot: downregulated ISC genes (FDR<0.05, likelihood-ratio test, grey dots: non-DE genes. **E.** Bar plot shows frequency (*y*-axis) of IEC-types, as determined by unsupervised clustering (**Methods**), in 815 cells from *Foxp3*-DTR mice (*n*=4, white) *vs*. 2,572 cells from matched WT controls (*n*=5, grey) both treated with DT. Individual mice marked by points. Error bars: SEM. (** FDR<0.005, *** FDR <10^-5^ likelihood-ratio test). **F.** Expansion of tuft cells following T_reg_ depletion in *Foxp3*-DTR mice. IFA image of DCLK1^+^ (red) tuft cells in the epithelia of wild type (WT, left) and *Foxp3-*DTR mice (right) both treated with DT. Scale bar, 50μm.

